# Molecular determinants of chaperone interactions on MHC-I for folding and antigen repertoire selection

**DOI:** 10.1101/779777

**Authors:** Andrew C. McShan, Christine A. Devlin, Sarah A. Overall, Jihye Park, Jugmohit S. Toor, Danai Moschidi, David Flores-Solis, Hannah Choi, Sarvind Tripathi, Erik Procko, Nikolaos G. Sgourakis

## Abstract

The interplay between a highly polymorphic set of MHC-I alleles and molecular chaperones shapes the repertoire of peptide antigens displayed on the cell surface for T cell surveillance. Here, we demonstrate that the molecular chaperone TAPBPR associates with a broad range of partially folded MHC-I species inside the cell. Bimolecular fluorescence complementation and deep mutational scanning reveal that TAPBPR recognition is polarized towards one side of the peptide-binding groove, and depends on the formation of a conserved MHC-I disulfide epitope in the α_2_ domain. Conversely, thermodynamic measurements of TAPBPR binding for a representative set of properly conformed, peptide-loaded molecules suggest a narrower MHC-I specificity range. Using solution NMR, we find that the extent of dynamics at “hotspot” surfaces confers TAPBPR recognition of a sparsely populated MHC-I state attained through a global conformational change. Consistently, restriction of MHC-I groove plasticity through the introduction of a disulfide bond between the α_1_/α_2_ helices abrogates TAPBPR binding, both in solution and on a cellular membrane, while intracellular binding is tolerant of many destabilizing MHC-I substitutions. Our data support parallel TAPBPR functions of *i)* chaperoning unstable MHC-I molecules at early stages of their folding process, akin to a holdase with broad allele-specificity, and *ii)* editing the peptide cargo of properly conformed MHC-I molecules *en route* to the surface, which demonstrates a narrower specificity. Our results suggest that TAPBPR exploits localized structural adaptations, both near and distant to the peptide-binding groove, to selectively recognize discrete conformational states sampled by MHC-I alleles, towards editing Sithe repertoire of displayed antigens.

**Significance Statement:** The human population contains thousands of MHC-I alleles, showing a range of dependencies on molecular chaperones for loading of their peptide cargo, which are then displayed on the cell surface for T cell surveillance. Using the chaperone TAPBPR as a model, we combine deep mutagenesis with functional and biophysical data, especially solution NMR, to provide a complete view of the molecular determinants of chaperone recognition. Our data provide significant evidence that localized protein motions define the intrinsic ability of MHC-I molecules to interact with chaperones. The importance of MHC-I dynamics unifies all our findings, with broad recognition of conformationally unstable, nascent MHC-I molecules becoming restricted to a smaller set of MHC-I alleles that retain relevant dynamic motions in their folded state.

**Graphical Abstract:** 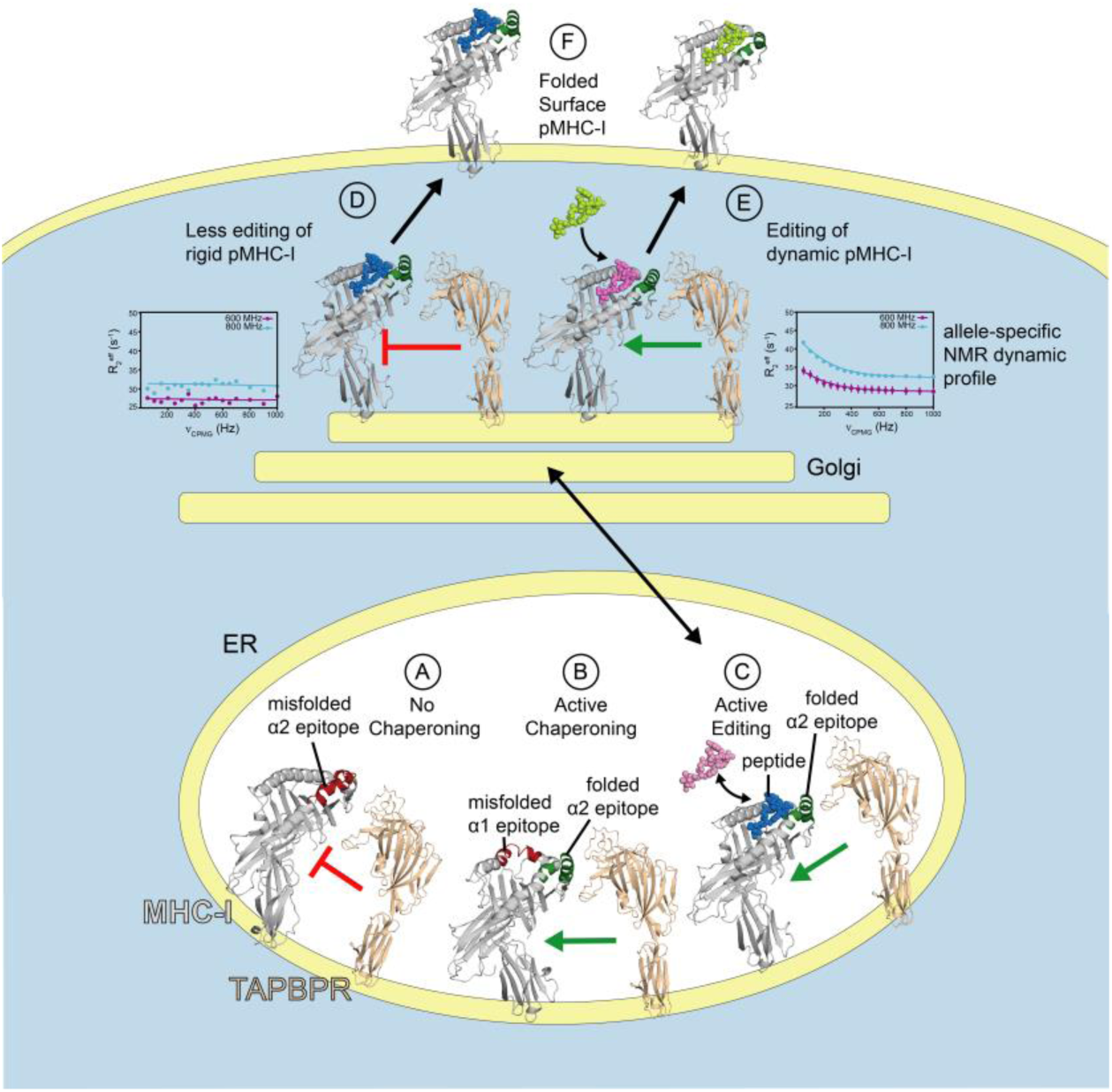

**Highlights:**

- Deep mutagenesis identifies a conformational disulfide-linked epitope as the main requirement for association of nascent MHC-I molecules with the TAPBPR chaperone
- Analysis of μs-ms timescale conformational dynamics by methyl NMR reveals allele-specific profiles at the TAPBPR interaction surfaces of peptide-loaded MHC-I molecules
- μs-ms dynamics dictate the specificity of TAPBPR interactions for different MHC-I alleles through the sampling of a minor, “excited state” conformation
- Restriction of dynamics though an engineered disulfide bond abrogates interactions with TAPBPR, both in solution and on a cellular membrane

## Introduction

On the surface of nucleated cells of jawed vertebrates, class I major histocompatibility complex (MHC-I) molecules display a diverse set of 8 to 14 residue peptide antigens to CD8^+^ cytotoxic T-lymphocytes (*1*). This process provides a means for immune surveillance of the endogenous proteome to detect invading pathogens or developing tumors. MHC-I molecules are extremely polymorphic, with thousands of known human alleles, categorized in the HLA-A, -B and -C classes. Specific interactions with highly polymorphic pockets along the peptide binding groove (termed A-F pockets) define a repertoire of up to 10^4^ peptide antigens that can bind to each HLA protein (*1*). Proper folding of nascent MHC-I molecules and loading with high-affinity peptides requires association with an invariant light chain, β2-microglobulin (β2m), and is facilitated by dedicated molecular chaperones, tapasin, which is restricted within the peptide-loading complex (PLC), and the homologous, PLC-independent TAPBPR (TAP-binding protein related) (*2, 3*). Furthermore, through the catalytic enhancement of peptide association and dissociation within the MHC-I groove, chaperones can influence the selection of immunodominant antigens by promoting the exchange of low- and intermediate-affinity for high-affinity peptides (termed peptide editing) (*2, 3*). Further quality control and narrowing of the displayed antigen repertoire along the trafficking pathway is accomplished through the combined functions of TAPBPR and UDP-glucose:glycoprotein glucosyltransferase (UGGT) (*3–5*). Aberrant expression of MHC-I molecules and their chaperones has been associated with the onset of several immunological disorders (*6–9*). Mass spectrometry studies have provided a direct link between chaperone function and immune surveillance, by demonstrating differences in the repertoire of peptides displayed by MHC-I molecules on the surface of wild-type versus tapasin (or TAPBPR) knockout cells (*4, 5, 10, 11*).

The discovery that TAPBPR can function as a peptide exchange catalyst outside the peptide-loading complex and can maintain empty MHC-I molecules in a peptide-receptive conformation has opened a new window to study the peptide loading process in a range of detailed functional and mechanistic studies (*4, 12, 13*). Structural insights into static snapshots of MHC-I/chaperone complexes have been gleaned by X-ray crystallography (*14, 15*) and cryoEM (*16*). However, MHC-I molecules are inherently dynamic at multiple sites (*17, 18*) and transitions between different “open” and “closed” conformations enable peptide loading, as shown by molecular dynamics (MD) simulation (*19, 20*) and kinetic modeling (*21*). In recent work, we have used methyl-based NMR to provide direct insights into the dynamics of conserved MHC-I regions in the TAPBPR complex. Together, these studies have shown that chaperones *(i)* stabilize peptide-deficient MHC-I in a peptide-receptive conformation with a widened groove, *(ii)* eject suboptimal peptides from the MHC-I and *(iii)* dissociate upon binding of high-affinity peptides, due to structural adaptations of a conformational “latch” located on the α_2-1_ helix, near the F-pocket (*22*).

Improper function of the antigen processing and presentation pathway confers susceptibility to diseases in a manner that is highly dependent on the individual’s MHC-I haplotype and the disease-relevant immunodominant peptides (*23*). Several studies have provided key insights into the allelic and peptide preferences of chaperone interactions (*4, 12, 13, 23, 24*). With regards to peptide preferences, TAPBPR has been demonstrated to form high-affinity complexes with MHC-I molecules that are either peptide-deficient or loaded with suboptimal peptides (*4, 13*). However, it remains unclear how intrinsic dissociation of low-affinity peptides from the MHC-I groove, occurring at the milliseconds timescale, affects chaperone binding measurements *in vitro* (*25*). In experiments performed under non-equilibrium conditions, such as size exclusion chromatography (SEC), surface plasmon resonance, or measurements on the cell surface using fluorescence methods, peptide dissociation from the MHC-I groove over the course of the experiment could influence the apparent chaperone dissociation rate. Solution NMR allows the simultaneous tracking of MHC-I species of different peptide occupancies via their unique chemical shifts, under equilibrium conditions. Using NMR, we have demonstrated that a soluble form of TAPBPR can recognize MHC-I molecules loaded with high-affinity peptides, albeit with micromolar affinity relative to nanomolar affinity when the groove is empty or partially loaded with peptides (*15, 19*). With respect to the allelic dependence of chaperone interactions, some MHC-I alleles require tapasin for proper peptide loading, trafficking and cell surface display, while others can intrinsically load peptides in tapasin knockouts (*4, 13, 24, 26*). In an extreme case demonstrated by the HLA-B*44 alleles, a single amino acid polymorphism located in the MHC-I groove is sufficient to switch between chaperone-dependent and independent peptide loading, with the chaperone independent allele linked with susceptibility to ankylosing spondylitis (*27–29*). Similarly, TAPBPR expression exhibits a marked effect on the displayed repertoire of specific alleles (*4*). While structure-based modeling using the available co-crystal structures as templates can provide some clues for the molecular basis of MHC-I allelic dependence of chaperone interactions, MD simulations and hydrogen deuterium exchange experiments have further shown that chaperone-dependent and independent alleles exhibit widespread differences in their dynamic profiles (*30, 31*). In particular, these experiments have suggested that differences in the rate of sampling of an “open” MHC-I conformation for different alleles confers chaperone recognition (*32, 33*).

Despite these important insights, the molecular determinants that underpin interactions of MHC-I alleles with chaperones at different stages of the folding and peptide loading processes remain incompletely understood. Here, using the chaperone TAPBPR as a model, we apply a range of complementary functional and biophysical experiments on fusion protein probes and recombinant molecules in a native cellular context and in solution, respectively. Our results provide a comprehensive characterization of the sequence, structure and dynamic features that confer TAPBPR recognition. We find that TAPBPR interacts broadly with nascent MHC-I of varying sequences within cellular compartments, with a binding mode that is highly polarized towards one side of the peptide-binding groove, and may function as a “holdase” of misfolded molecules. However, interactions with properly conformed pMHC-I molecules towards editing of the peptide cargo are restricted to a limited set of alleles, where the dynamic sampling of a sparse minor-state conformation in solution is important. A role for MHC-I dynamics unifies these findings, with broad recognition of conformationally unstable nascent MHC-I becoming restricted to a smaller set of alleles that retain relevant dynamic motions in the folded and peptide-loaded state. Our results provide a molecular blueprint for the design of MHC-I molecules with improved antigen processing and presentation properties, through the engineering of conformational species across the entire folding landscape.

## Results

### TAPBPR associates with a broad range of MHC-I alleles inside the cell

TAPBPR has been reported to exhibit a narrow MHC-I allele specificity (*4, 13*). However, it remains undetermined whether the allelic preferences of TAPBPR manifest in a similar manner for nascent, empty MHC-I or properly conformed, peptide-loaded MHC-I molecules. To explore these concepts further, we first investigated the propensity for TAPBPR to interact with MHC-I molecules composed of different heavy chain sequences (mouse H2 or human HLA) in a native cellular environment. We adapted established bimolecular fluorescence complementation (BiFC) assays in human Expi293F cells transfected using gene constructs containing TAPBPR fused to the C-terminal half (VC) of split fluorescent Venus, and one of a panel of heavy chain MHC-I fused to the N-terminal half (VN) (*34*). Here, when TAPBPR-VC comes into proximity with MHC-I-VN as the result of protein-protein interactions inside the cell, an active yellow fluorescent protein can fold resulting in high BiFC signal which can be readily detected by flow cytometry. As a negative control, we used the unrelated protein CXCR4-VN (C-X-C chemokine receptor type 4), which is not expected to interact with TAPBPR-VC. After gating for cells with similar expression levels based on intracellular staining of N-terminal epitope tags (**Fig. S1A-C**), we observe significant BiFC signal across all H2 and HLA alleles tested, suggesting that a broad range of MHC-I alleles with diverse sequences have the potential to interact with TAPBPR in a native cellular environment (**Fig. S1D**). Background BiFC signal for CXCR4 from non-specific protein associations is substantially lower. We note that gating based on expression levels did not change observed trends, but rather highlighted differences by excluding high-expressing cells with saturated BiFC signal (**Fig. S1A, B**). Overexpression of TAPBPR in this assay also promotes intracellular retention of MHC-I, consistent with previous reports (*12*). Unlike the negative control CXCR4, the MHC-I molecules tested here exhibit severely reduced cell surface expression in the presence of TAPBPR (**Fig. S1E**).

Spontaneous and non-specific self-assembly of the Venus halves due to protein overexpression can result in high background and low signal-to-noise, leading to false-positive BiFC signals for some protein-protein interactions (*34*). Thus, we co-expressed and co-immunoprecipitated TAPBPR with MHC-I in the absence of VN and VC fusions. Unlike the CXCR4 control, all the MHC-I alleles bound TAPBPR), thereby independently confirming the BiFC results using a different experimental method (**Fig. S2**). Together, these data suggest that TAPBPR associates with a broad range of MHC-I alleles inside the cell, potentially through partially folded, empty MHC-I conformations within the endoplasmic reticulum (ER) or Golgi.

**Fig. 2.**
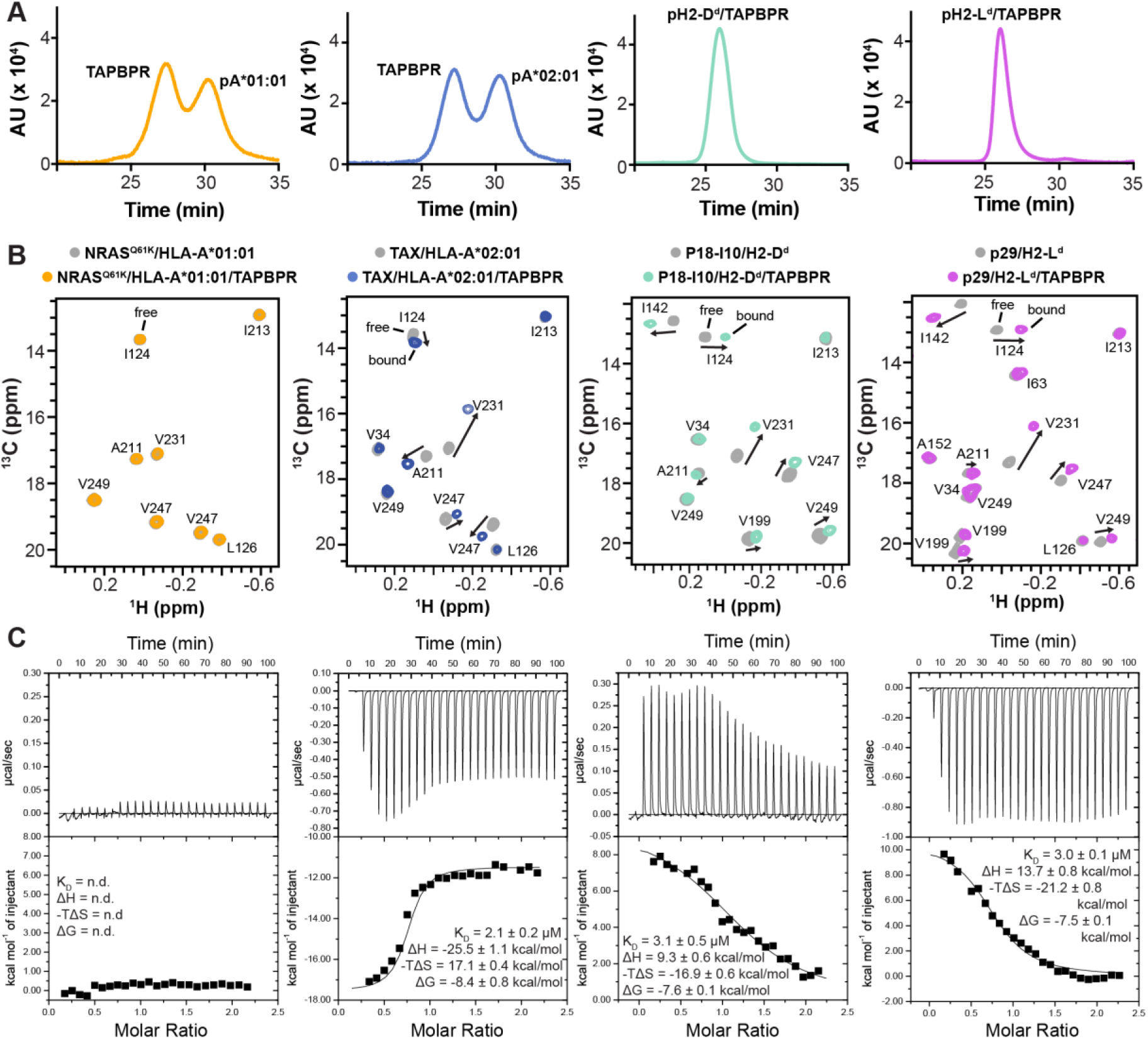
TAPBPR interactions with peptide-loaded MHC-I molecules are allele-dependent. **(A)** SEC traces of 1:1 molar ratio of TAPBPR with different pMHC-I heavy chains, each refolded with human β2m and a full-length peptide. **(B)** Representative 2D ^1^H-^13^C HMQC spectra of heavy chain ^13^C AILV methyl labeled pMHC-I at 105 μM without TAPBPR (gray) and in the presence of three-fold molar excess TAPBPR. From left to right: NRAS^Q61K^/HLA-A*01:01/hβ2m (orange), TAX/HLA-A*02:01/hβ2m (blue), P18-I10/H2-D^d^/hβ2m (green), and NIH/H2-L^d^/hβ2m (purple). Experiments were performed at a ^1^H NMR field of 800 MHz at 25°C. Arrows indicate significant chemical shift changes in MHC-I heavy chain NMR resonances upon TAPBPR binding. **(C)** Representative ITC data titrating ∼100 to 150 μM pMHC-I (from left to right: NRAS^Q61K^/HLA-A*01:01, TAX/HLA-A*02:01, P18-I10/H2-D^d^ and p29/H2-L^d^) into a sample containing 12 μM TAPBPR and 1 mM peptide. Black lines are the fits of the isotherm. Fitted values for K_D_, ΔH, -TΔS, and ΔG were determined using a one-site binding model. n.d., not determined. Standard errors were determined from experimental replicates (n = 2).

### TAPBPR recognizes a partially folded epitope on MHC-I molecules with broad specificity

Deep mutagenesis, in which directed evolution of a diverse library of sequences is tracked by next generation sequencing, has emerged as a powerful tool for interrogating how protein amino acid sequence dictates structure and function in living cells (*35*). The relative activities of thousands of mutations can be assessed simultaneously to reveal critical features for protein stability or ligand binding. To identify important features of the human MHC-I heavy chain required for TAPBPR association within cells, we generated a single site-saturation mutagenesis (SSM) library on HLA-A*02:01. We chose to focus on HLA-A*02:01 as it is a common human allele that associates with TAPBPR *in vitro* with high affinity and presents several important autoimmune, viral and cancer epitopes (*4, 13*). To improve sampling of individual mutations and thereby increase data quality, library diversity was limited by restricting mutational scanning to the MHC-I α_1_/α_2_ “platform” domain (residues 2 to 181), which mediates both peptide and TAPBPR binding (*14, 15*). Using two complementary fluorescence-based selections, we investigated the effects of single site mutations throughout the groove on *i*) HLA-A*02:01 cell surface expression, to reveal how particular mutations influence MHC-I stability, and *ii*) HLA-A*02:01/TAPBPR recognition, to identify structural elements that are important for chaperone recognition. There were concerns that high BiFC signal may simply report on assembly with MHC-I-VN mutants that are retained in the same intracellular compartments as TAPBPR-VC (see above). To resolve this potential issue, we used cells stably expressing a variant called TAPBPR-TM, in which the native transmembrane helix and cytosolic tail are replaced with a generic transmembrane domain. Confocal microscopy (**Fig. S3A**) and flow cytometry (**Fig. S3B**) reveal that TAPBPR-TM localizes to both the Golgi and plasma membrane, and therefore reports on interactions regardless of whether MHC-I mutants traffic to the surface or not. Following transfection of the HLA-A*02:01 SSM library and fluorescence activated cell sorting (FACS) for MHC-I surface expression in wild type cells (**Fig. S3C**) or MHC-I/TAPBPR BiFC in TAPBPR-TM-VC expressing cells (**Fig. S3D**), the relative effects of nearly all single amino acid substitutions in the MHC-I groove are determined based on their enrichment or depletion (**Fig. S4A, B)**. Enrichment ratios of specific mutations and conservation scores for residue positions have excellent agreement between independent replicates, providing high confidence to the mutational landscapes (**Fig. S4C-G**).

We find that surface expression of HLA-A*02:01 imposes strict sequence constraints throughout the peptide binding groove, consistent with the preferential trafficking of stable, peptide-loaded MHC-I to the plasma membrane (**Fig. 1A, B and Fig. S4A**) (*36, 37*). Not only are the residues comprising the MHC-I pockets that anchor the peptide tightly conserved for surface expression (residues 5, 7, 59, 63, 66, 99, 159, 163, 167, 171 for the A-pocket and 77, 80, 81, 84, 116, 123, 143, 146, 147 for the F-pocket), but so are residues within the core of the MHC-I groove, and the underlying surface contacting β2m and the α_3_ domain (**Fig. 1A-C and Fig. S4A)**. Proper pMHC-I folding and stability in human cells, which express endogenous tapasin, is therefore intolerant of most single amino acid substitutions. In an interesting exception to this trend, sites on the α1 helix (MHC-I residues 67-75) that do not participate in direct interactions with either the bound peptide or TAPBPR, but would be engaged by T cell receptors, are more amenable to mutation (**Fig. 1A and Fig. S4A**). Together, these data reveal that conserved MHC-I sites important for surface expression are dispersed throughout the groove, and that the most stringent sequence constraints are contained within the peptide anchoring pockets, consistent with the established trend that MHC-I surface expression is tightly coupled to proper peptide loading (*36, 37*).

**Fig. 1.**
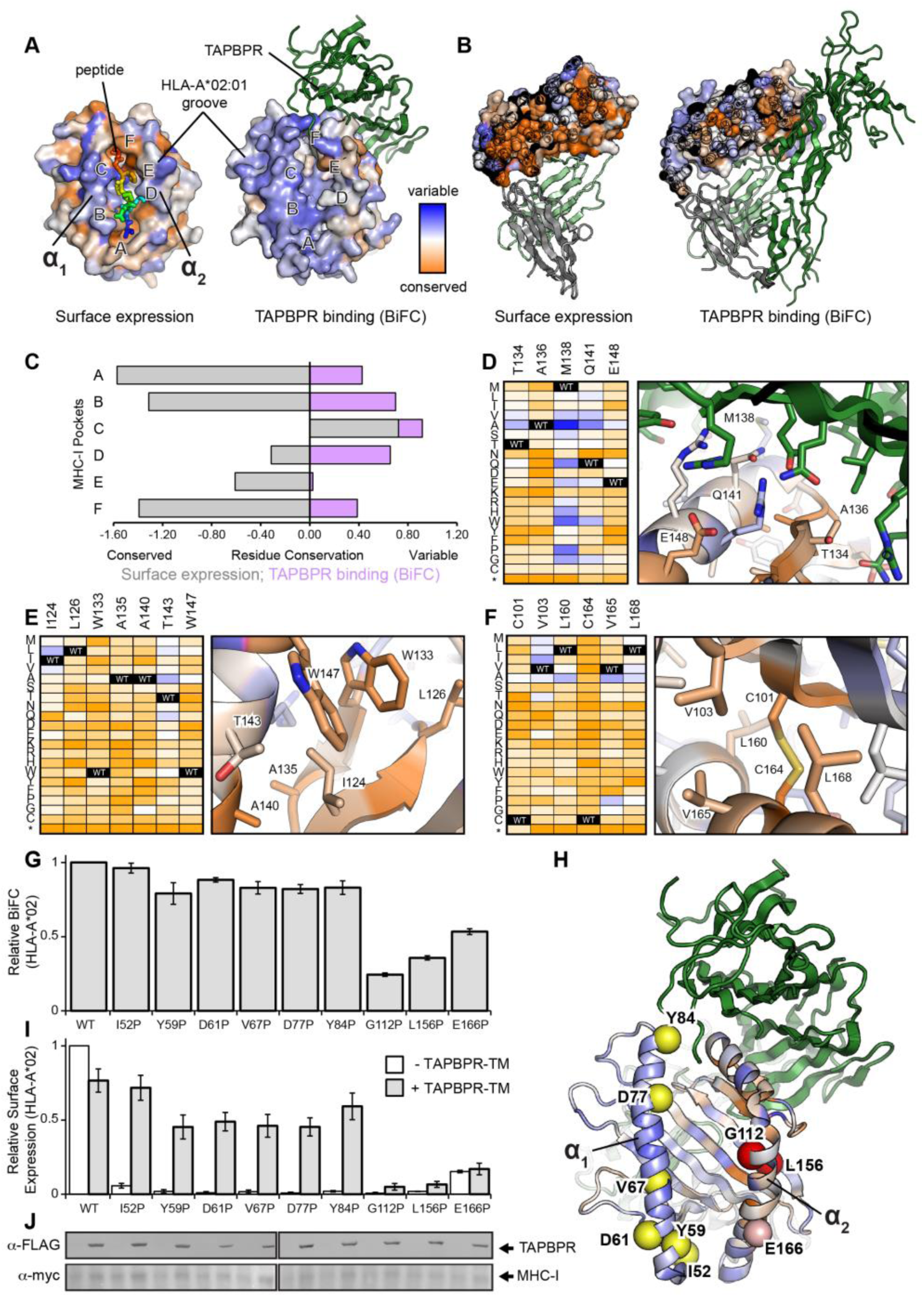
TAPBPR recognizes a local conformation of the HLA-A*02:01 α_2_ domain. **(A)** Sequence conservation from deep mutagenesis mapped to the surface of HLA-A*02:01. Conserved residues are colored in dark orange, while residues exhibiting mutational tolerance in pale and blue. Residue conservation for HLA-A*02:01 surface expression is shown on the structure of the free pMHC-I molecule (PDB ID 1HHJ), with bound peptide colored from blue (P1) to red (P9). HLA-A*02:01 conservation for TAPBPR (dark green) binding is plotted on the structure of the empty MHC-I, in complex with TAPBPR (PDB ID 5WER). The location of the A- to F-pockets of the MHC-I groove are noted. **(B)** Cross sections through the core of the α_2_ domain, colored by conservation as in panel A. TAPBPR is dark green, β2m is pale green, and the α_3_ domain is grey. **(C)** Conservation and variation of residues in the MHC-I pockets for surface expression (gray) or TAPBPR binding by BiFC (purple). **(D-F)** Heat maps of mutation log_2_ enrichment ratios for HLA-A*02:01/TAPBPR BiFC (depleted mutations are orange, enriched mutations are dark blue) shown alongside modeled structures of the (D) α_2_ domain/TAPBPR interface, (E) the hydrophobic core between the α_2-1_ helix and β-sheet, (F) and the core between the α_2-2_ helix and β-sheet. **(G)** Relative BiFC signal between TAPBPR-TM-VC and MHC-I-VN for different proline substitutions in either the α_1_ or α_2_ helices, or β-sheet underlying the α_2_ helix. **(H)** Location of proline substitutions on the MHC-I groove mapped onto the TAPBPR complex structure (PDB ID 5WER). **(I)** Relative surface expression for the different MHC-I proline substitutions in the presence or absence of TAPBPR-TM. **(J)** Immunoblots comparing total expression levels for TAPBPR-TM (α-FLAG) and HLA-A*02:01 (α-myc) constructs. Lanes are aligned with graphs above.

While surface expression imposes strict sequence constraints throughout the MHC-I groove, we find that TAPBPR association persists following many mutations, and is especially tolerant of mutations to the 3_10_ and α_1_ helices that demarcate one side of the peptide binding groove (**Fig. 1A-C)**. This also includes tolerance to many mutations at positions for which crystallographically defined intermolecular contacts are observed in the homologous mouse H2-D^d^/TAPBPR complex (*15*), such as HLA-A*02:01 residues 83, 84, 127 and 144 (**Fig. 1A, B and Fig. S4B**). Other HLA-A*02:01 residues that are buried in the homologous H2-D^d^/TAPBPR interface, such as 134, 135, 136, and 148, are under tight sequence conservation (**Fig. 1D**), revealing that only a subset of contacts at the MHC-I/TAPBPR interface are malleable to mutation. While HLA-A*02:01 residues comprising the A-, B, and F-pockets of the groove are conserved for surface expression, they tolerate variability for TAPBPR binding (**Fig. 1C**), consistent with TAPBPR engaging HLA-A*02:01 mutants that are defective for peptide loading. There are two prominent buried clusters of conserved MHC-I residues in the mutational landscape. The first cluster contains HLA-A*02:01 residues from the region 124-147, located within and adjacent to the E- and F-pockets **(Fig. 1D, E**). A second, smaller cluster of MHC-I residues conserved for TAPBPR binding mediates contacts between the α_2-2_ helix and β-sheet around the conserved C101-C164 disulfide within the A-pocket (**Fig. 1F**). We find that mutation of the C101-C164 disulfide abrogates interaction with TAPBPR, likely because disulfide bond formation contributes to stabilization of the α_2_ domain (*38*). Elsewhere, HLA-A*02:01/TAPBPR BiFC shows intermediate mutational tolerance at sites of the MHC-I groove contacting the β2m/α_3_ domains, while the core between the α_1_ helix and β-sheet has very high mutational tolerance (**Fig. S4B)**. Thus, in contrast to the strict sequence conservation of MHC-I sites dispersed throughout the groove for surface expression, sites conserved for TAPBPR association are highly polarized towards the α_2_ domain of the groove, suggesting that TAPBPR has the potential to recognize nascent MHC-I molecules with broad sequence specificity, at least when single amino acid substitutions are considered.

### TAPBPR recognition of nascent species is polarized towards one side of the MHC-I groove

We hypothesized that TAPBPR can associate with partially folded or misfolded MHC-I molecules, with an interaction mode that is dependent upon the formation of a local conformational epitope adjacent to the α_2_ helix. Such MHC-I species likely form intracellularly during the processing of nascent, peptide-deficient molecules. To investigate this, we created a panel of proline substitutions along either the α_1_ or α_2_ helices of HLA-A*02:01 groove. Incorporation of a proline is expected to destabilize secondary structure, especially helices (*39, 40*). Destabilizing proline substitutions within the α_1_ helix (I52P, Y59P, D61P, V67P, D77P, Y84P) do not abrogate BiFC signal with TAPBPR-TM compared to wild-type HLA-A*02:01, while those within or adjacent to the α_2_ helix (G112P, L156P, E166P) severely reduce BiFC signal with TAPBPR-TM (**Fig. 1G, H**). We simultaneously examined MHC-I surface localization in the transfected cells. Ordinarily, destabilized MHC-I proline mutants would not escape intracellular quality control to reach the plasma membrane, but stable association with TAPBPR-TM, which traffics to the cell surface, is expected to rescue MHC-I surface localization. Indeed, we find that proline substitutions within the α_1_ helix of the groove are able to reach the cell surface only in the presence of TAPBPR-TM, while those within the α_2_ helix are retained intracellularly in the absence and presence of TAPBPR-TM (**Fig. 1I**). Importantly, expression of all the mutants remain at near wild type levels, and hence observed differences cannot be explained by a simple loss of expression (**Fig. 1J**). Thus, while destabilization of the HLA-A*02:01 α_1_ helix induces a misfolded, peptide-deficient state, these conformations do not compromise the TAPBPR recognition epitope and a stable complex can form (**Fig. 1H**). This is in contrast with proline substitutions within or near to the α_2_ helix, which disrupt the local MHC-I structure recognized by TAPBPR (**Fig. 1H**).

### TAPBPR recognizes folded pMHC-I using a conserved binding mode of narrow specificity

To characterize TAPBPR recognition of purified pMHC-I molecules in solution, we focused on a representative set corresponding to two mouse - H2-D^d^ and H2-L^d^, and two human - HLA-A*01:01 and HLA-A*02:01 alleles, displaying between 70 to 85% similarity at the sequence level (**Fig. S5**). Each MHC-I heavy chain was expressed in *E. coli*, and refolded together with a high-affinity peptide and human β2m (hβ2m) (*13, 19*). We utilized well-characterized epitopes: the p29 peptide (YPNVNIHNF) for H2-L^d^ (*41*), the HIV gp120 P18-I10 peptide (RGPGRAFVTI) for H2-D^d^ (*42*), the neuroblastoma related NRAS^Q61K^ peptide (ILDTAGKEEY) for HLA-A*01:01 (*43*), and the HTLV-1 TAX peptide (LLFGYPVYV) for HLA-A*02:01 (*44*). Using a differential scanning fluorimetry assay (*45*), we further confirmed that all purified samples displayed thermal stabilities that were characteristic of properly conformed, peptide-bound molecular species with T_m_ values ranging from 51 to 63 °C (**Table S1**). Using a SEC assay, we observed formation of pMHC-I/TAPBPR complexes for both mouse pMHC-I (P18-I10/H2-D^d^/hβ2m and NIH/H2-L^d^/hβ2m), but neither of the human pMHC-I (NRAS^Q61K^/HLA-A*01:01/hβ2m and TAX/HLA-A*02:01/hβ2m) (**Fig. 2A**). A potential drawback of SEC is that measurements carried out under non-equilibrium conditions may favor complex and/or peptide dissociation, depending on kinetic off-rates (*46*). Thus, we also monitored TAPBPR binding under equilibrium conditions using solution NMR. Here, we used isotopically AILV-methyl labelled (Ala ^13^Cβ; Ile ^13^Cδ1; Leu ^13^Cδ1/^13^Cδ2; Val ^13^Cγ1/^13^Cγ2) pMHC-I (where the hβ2m subunit and peptide are at natural isotopic abundance), which allows us to quantify specific chemical shift changes upon formation of the 87 kDa pMHC-I/TAPBPR complex (*19*). NMR can also detect the presence of peptide-deficient complex in the sample, through the observation of unique chemical shifts of MHC-I methyl groups of residues in the peptide-binding groove (*19*). Towards this goal, we obtained complete stereospecific assignment for the heavy chain AILV methyl groups of each pMHC-I system in our set (*19, 47*). Simultaneously, to obtain precise thermodynamic parameters describing the binding process, we performed isothermal calorimetry titration (ITC) experiments (*48*). To ensure that the complex formed with TAPBPR contained exclusively peptide-bound MHC-I (rather than empty molecules), all ITC experiments were performed in the presence of 10-fold molar excess of free peptide relative to the pMHC-I (see Materials & Methods).

Both NMR and ITC experiments confirm that peptide-bound TAX/HLA-A*02:01/hβ2m, P18-I10/H2-D^d^/hβ2m and NIH/H2-L^d^/hβ2m form high-affinity complexes with TAPBPR *in vitro* (**Fig. 2B, C and Fig. S6**). In NMR experiments, tight complex formation is highlighted by the slow-exchange chemical shift changes of resonances corresponding to methyls groups of conserved heavy chain residues, such as I124, A211, V231 and V247 (**Fig. 2B and Fig. S6**), suggesting a similar TAPBPR binding mode. Peptide release from the pMHC-I/TAPBPR complex to form empty complex was not detected under the NMR sample conditions (**Fig. 2B**). Whereas the binding footprint of TAPBPR probed by methyl NMR is highly conserved among the three interacting pMHC-I molecules, and is further consistent with the published X-ray structures (*14, 15*), our ITC data reveal that the same complex structure can be attained via distinct thermodynamic processes. Specifically, HLA-A*02 exhibits exothermic binding behavior, in contrast to the endothermic association observed with H2-D^d^ and H2-L^d^. While the enthalpic (ΔH) and entropic (-T*ΔS) contributions may differ between HLA-A*02:01 and H2-D^d^/H2-L^d^, the resulting binding free energy (ΔG) values are very similar (**Fig. 2C**). Fitting of the sigmoidal isotherms using a one-site interaction model, yields very similar apparent dissociation constants of 2.1 μM (TAX/HLA*A*02:01/ hβ2m), 3.1 μM (P18-I10/H2-D^d^/hβ2m) and 3.0 μM (NIH/H2-L^d^/hβ2m) (**Fig. 2C**). Notably, NRAS^Q61K^/HLA-A*01:01/hβ2m does not interact with TAPBPR by either SEC, NMR or ITC, up to millimolar-range concentrations (**Fig. 2A-C and Fig. S6**). Together, our biophysical data reveal that relative to the broad specificity exhibited towards MHC-I molecules of different sequence features inside the cell (**Fig. S1 and Fig. S2**), TAPBPR recognizes properly conformed, peptide-loaded MHC-I molecules with a narrow specificity in solution, even when comparing pMHC-I complexes that are very similar in terms of overall fold, amino acid composition and thermal stability.

**Fig. 6.**
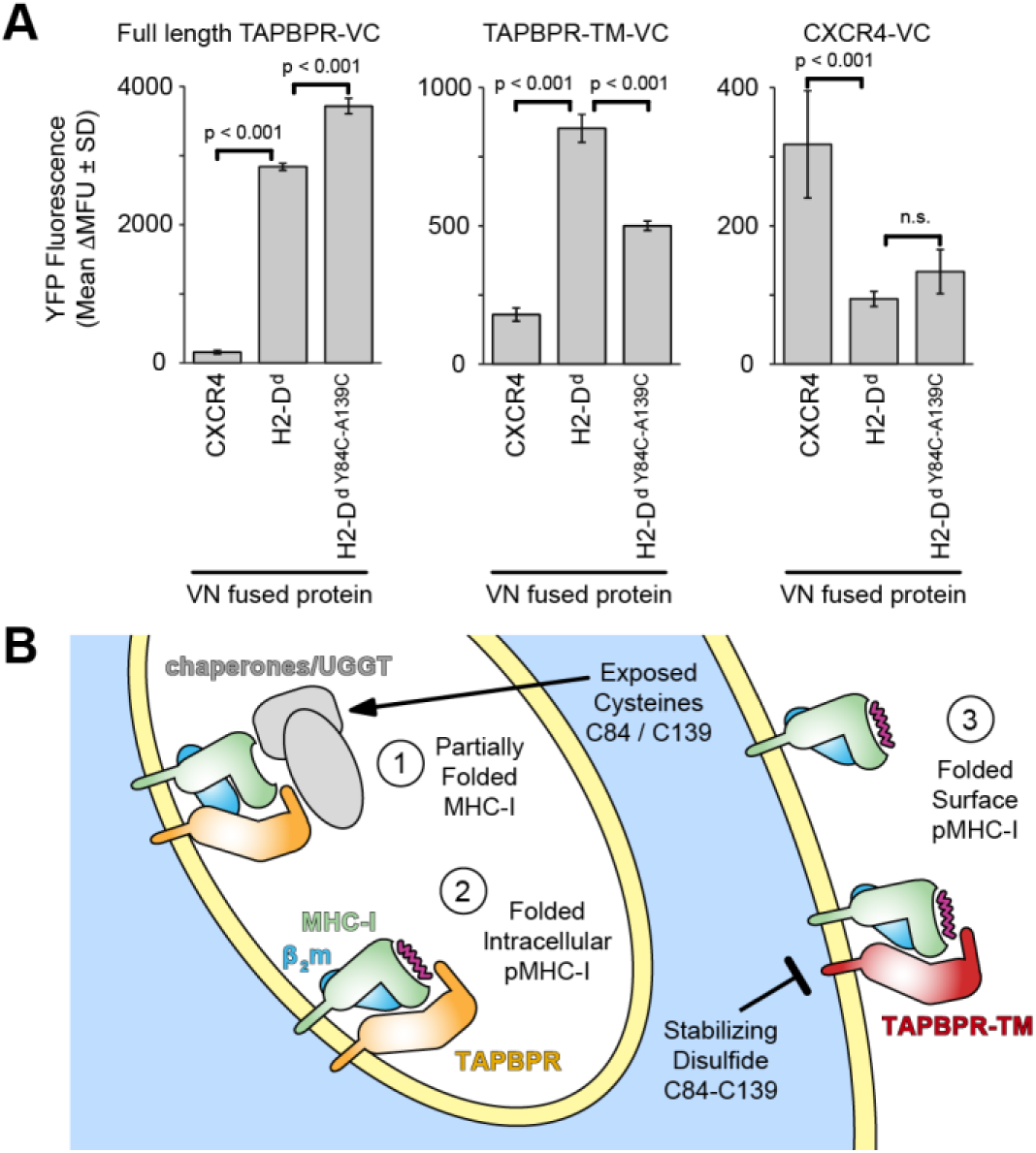
Restriction of MHC-I dynamics diminishes TAPBPR association on a cell membrane. **(A)** Cells were co-transfected with VC-fused FLAG-TAPBPR (left), FLAG-TAPBPR-TM (middle) or FLAG-CXCR4 (right), and with VN-fused myc-H2-D^d^ or myc-CXCR4. Yellow fluorescence (shown as average ΔMean Fluorescence Units ± SD, n = 4) was measured after gating for equivalent expression levels. P-vales are calculated from two-tailed Student’s t test. **(B)** Nascent MHC-I (step 1) within intracellular compartments associates with quality control machinery. The presence of additional, partially reduced cysteines is hypothesized to enhance chaperone associations, including direct or indirect recruitment of TAPBPR. As MHC-I completes its folding, it may transiently interact with TAPBPR (step 2) before mature pMHC-I traffics to the plasma membrane (step 3). The stabilizing disulfide diminishes direct interactions between folded pMHC-I molecules and TAPBPR.

### pMHC-I molecules exhibit allele-specific conformational dynamics

Conformational dynamics have been implicated in several aspects of MHC-I function, including peptide loading, T cell receptor triggering, and chaperone recognition (*2, 17, 49*). Solution NMR allows the quantitative measurement of dynamics across biologically relevant timescales ranging from psec-nsec (side-chain and loop motions), μs-ms (minor domain movements) to seconds (major domain reorientation) (*50*). Previous NMR measurements of psec-nsec dynamics in pMHC-I molecules did not uncover molecular flexibility other than in flexible loop regions (*19, 51*). In contrast, MHC-I regions involved in peptide and chaperone recognition exhibit μs-ms exchange between minor conformational states (*19, 52*). We have previously established the use of methyl groups as highly sensitive NMR probes with widespread coverage of the entire MHC-I structure to characterize unchaperoned and chaperoned MHC-I complexes (*19*).

Here, we have adopted a similar approach to quantitatively compare and contrast conformational dynamics in our representative set of pMHC-I molecules of different allelic composition, towards uncovering differences that could explain variation trends in TAPBPR recognition. We examined dynamics of AILV methyl groups of unchaperoned pMHC-I (where the heavy chain is isotopically labeled, while hβ2m and the peptide are at natural isotopic abundance) using ^13^C single-quantum Carr-Purcell-Meiboom-Gill (^13^C-SQ CPMG) relaxation dispersion NMR experiments (*52, 53*). In these experiments, methyl groups undergoing conformational exchange at the μs-ms timescale exhibit marked changes in the measured effective relaxation rate (R_2_^eff^) as a function of the frequency of the refocusing pulses (ν_CPMG_) (*50, 53*). For each pMHC-I, we acquired ^13^C-SQ CPMG relaxation dispersion data at 25°C at two NMR field strengths (600 MHz/14.1 Tesla and 800 MHz/18.8 Tesla). All AILV methyl groups with observable dynamics (R_ex_ > 1 s^−1^) were fit to a two-state exchange model (**Fig. 3A**). This procedure allowed us to extract quantitative NMR parameters that report on conformational exchange, including the exchange rate constant (k_ex_ = k_AB_ + k_BA_), populations of the major and minor states (p_A_ and p_B_) and absolute values of the ^13^C chemical shift differences between the minor and the major state (|Δω|) (*50, 53*). CPMG experiments were performed in the presence of three-fold molar excess of peptide to minimize the population of aggregation-prone empty MHC-I, as the half-lives of moderate to high-affinity peptides have been reported in the 0.3 to 28 hour range (*54–56*). Specifically, we found no significant difference in the CPMG profiles and derived parameters in experiments performed in the absence or presence of excess TAX peptide (**Fig. S7**), suggesting that the excited state observed in our CPMG experiments does not correspond to a transition between the peptide-bound and peptide-free form, but rather to a conformational change occurring while the peptide remains bound in the MHC-I groove.

**Fig. 3.**
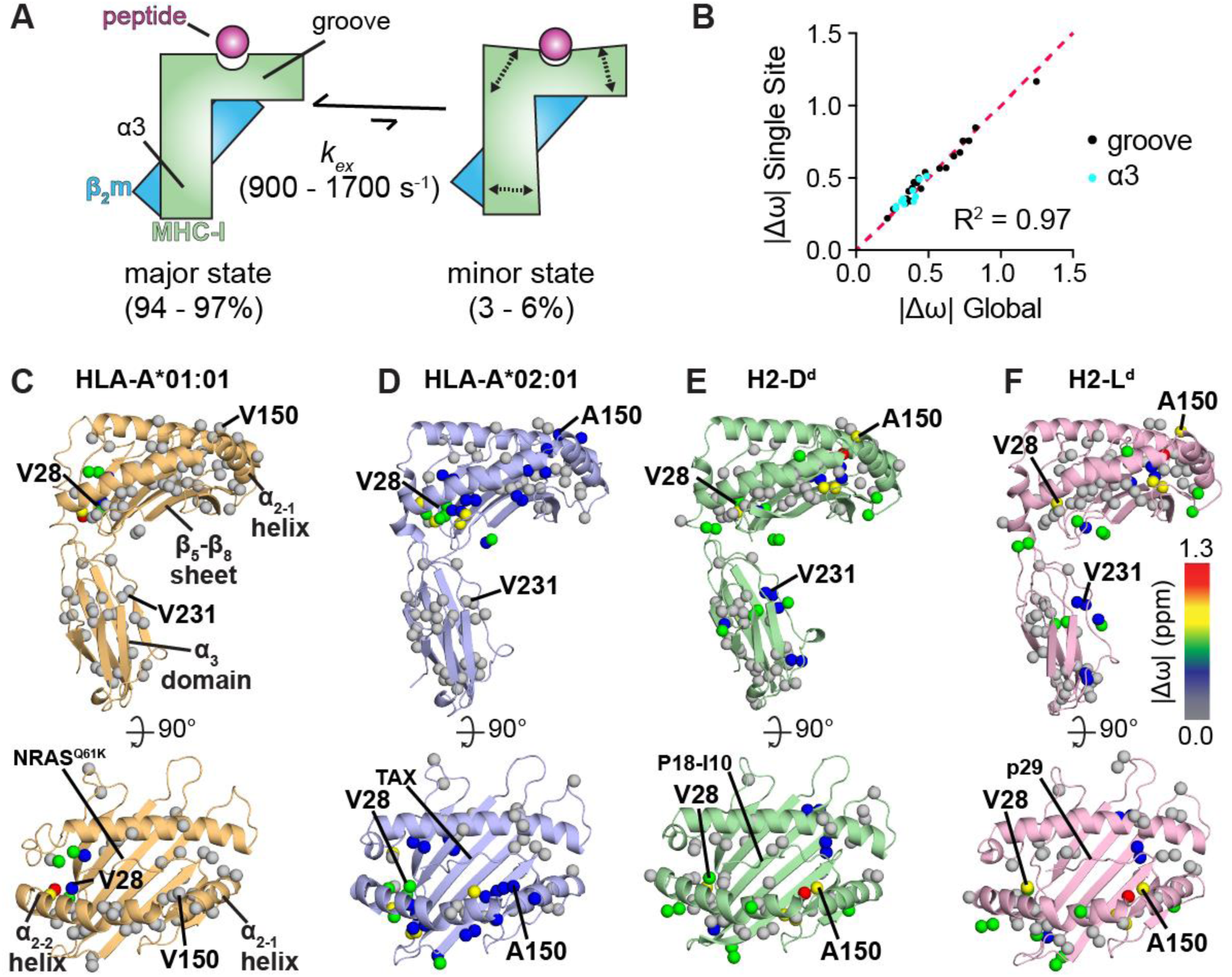
Dynamic profiles of different pMHC-I molecules probed by methyl NMR. **(A)** Summary of microsecond-milliseconds timescale conformational exchange of peptide-bound MHC-I molecules between a major, ground state and a minor, excited state. Experimentally determined populations of the major and minor states and the exchange rate (k_ex_) are noted in Table S2. **(B).** Comparison of |Δω| values obtained from a global fit of CPMG data for all sites together (*x*-axis) or independent fits of each site (*y*-axis), shown for methyl groups in the groove (black) or α_3_ domain (cyan). **(C-F)** The sites that participate in the global conformational exchange process are represented as spheres on the structure of each pMHC-I viewed from the side (top) or above the groove (bottom). hβ2m is omitted for clarity**. (C)** NRAS^Q61K^/HLA-A*01:01/hβ2m (PDB ID 6MPP) – orange, **(D)** TAX/HLA-A*02:01/hβ2m (PDB ID 1DUZ) (*44*) – blue, **(E)** P18-I10/H2-D^d^/hβ2m (PDB ID 3ECB) (*42*) – green and **(F)** NIH/H2-L^d^/hβ2m (PDB ID 1LD9) (*41*) – purple). Methyl groups undergoing dispersion are shown as spheres and color-coded based on |Δω| values obtained from a global fit of each pMHC-I. Methyl sites of increased dynamics are shown with warmer colors.

**Fig. 7.**
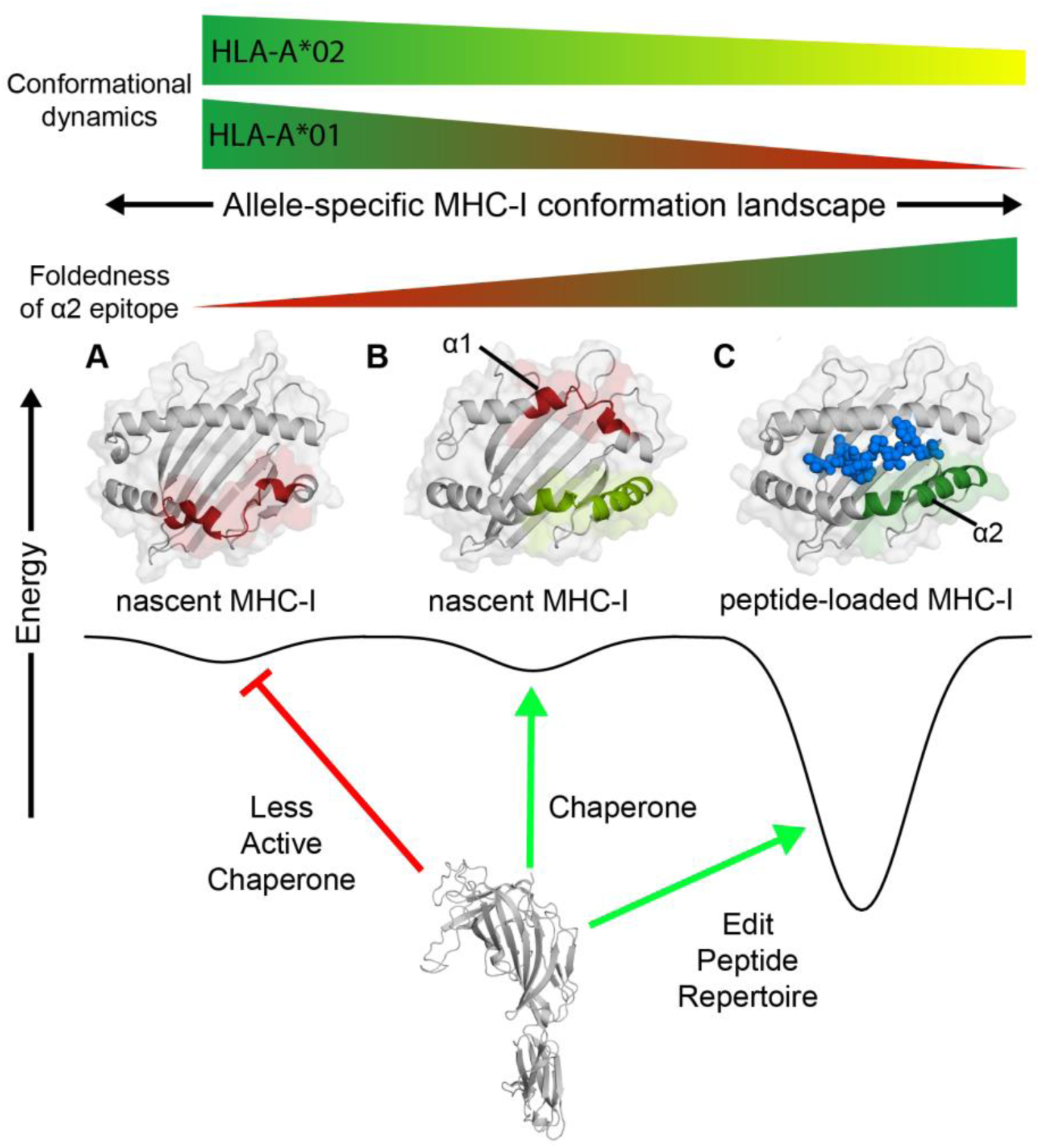
Chaperone recognition of a dynamic MHC-I conformational landscape. Conceptual example of the interaction between the chaperone TAPBPR and different MHC-I conformations of varying energetic and structural features. The vertical axis is free energy and the horizontal axis represents the conformational landscape of the MHC-I, which is influenced by specific polymorphisms in the MHC-I groove (α_1_/α_2_) and α_3_ domains. **(A)** TAPBPR does not associate with an MHC-I state comprising a misfolded α_2_ domain (red). **(B)** In the holdase function, TAPBPR interacts with nascent MHC-I conformations consisting of a folded α_2_ domain (light green) with an oxidized disulfide bond between the conserved Cys 101/164, even if the α_1_ domain remains in a misfolded state (red). **(C)** As a peptide editor, TAPBPR recognizes properly conformed molecules loaded with peptides towards exchange of the bound peptide cargo, for those alleles that exhibit μs-ms time scale conformational dynamics at the α_2-1_ helix (green). The peptide in the properly conformed pMHC-I state is shown as blue spheres.

We observed conformational exchange at pMHC-I sites important for function, including the α_2-1_ helix, the β_5_-β_8_ sheet forming the floor of the groove, the A-pocket and α_3_ domain. Notably, all CPMG profiles of methyl resonances showing conformational exchange could be individually fit to a model yielding the same population and kinetic parameters (**Fig. 3B**), suggesting the presence of a global conformational change connecting disparate sites on the MHC-I structure. However, the extent of dynamics within the different MHC-I domains was allele-dependent. While all pMHC-I molecules displayed A-pocket dynamics (**Fig. S8A** and **Fig. 3C-F,** residue 28), α_2-1_ helix dynamics were only observed in the three pMHC-I molecules recognized by TAPBPR (**Fig. S8B** and **Fig. 3C-F,** residue 150). In addition, μs-ms conformational exchange in the α_3_ domain was only observed for pMHC-I molecules prepared with mouse heavy chains H2-D^d^ or H2-L^d^ (**Fig. S8C** and **Fig. 3C-F**, residue 231). In all molecules, methyl groups span the entire length of the MHC-I groove and α_3_ domain, thus a lack of observed dynamics is not due to incomplete probe coverag**e (Fig. S9)**. Following a global fitting procedure of all methyl sites in each molecule, we obtained *k_ex_* values ranging from 976 to 1700 s^−1^ and *p_B_* values between 3.2 to 5.4% (**Table S2**). Fitted |Δω| values range between 0.2 and 1.3 ppm (**Fig. 3D-F** and **Table S2**). In summary, our CPMG data revealed unique allele-specific, dynamic profiles of unchaperoned pMHC-I molecules, for which differences were more pronounced at functionally relevant sites located within the α_2_ helix and α_3_ domain.

### A minor conformational state underpins TAPBPR recognition of different MHC-I alleles

To quantify and confirm that TAPBPR utilizes a conserved overall binding mode to recognize pMHC-I molecules of different allelic compositions, we performed a chemical shift deviation (CSD) analysis for complexes prepared using labelled HLA-A*02:01 and H2-L^d^ heavy chains, in addition to our previous characterization of H2-D^d^ (*19*). Here, the directly observed chemical shifts of the TAPBPR-bound state serve as unique identifiers of the induced local conformational changes, relative to the free pMHC-I state, for each residue. Using our established AILV methyl assignments for each system, the measured CSDs reveal that the same surfaces spanning multiple MHC-I domains on peptide-loaded H2-D^d^, H2-L^d^ and HLA-A*02:01 are engaged to form a tight complex with TAPBPR consistent with the peptide-deficient MHC-I/TAPBPR crystal structures (*14, 15, 19*) (**Fig. 4A, B**). Specifically, the NMR resonances corresponding to equivalent methyl probes in the α_1_ helix (residues 28-76), the β_5_-β_8_ strands (residues 104-130), the α_2-1_ helix (residues 139-158) and α_3_ domain (residues 230-251) of the heavy chain show significant CSDs upon TAPBPR binding across the three pMHC-I molecules analyzed (**Fig. 4A, B**).

**Fig. 4.**
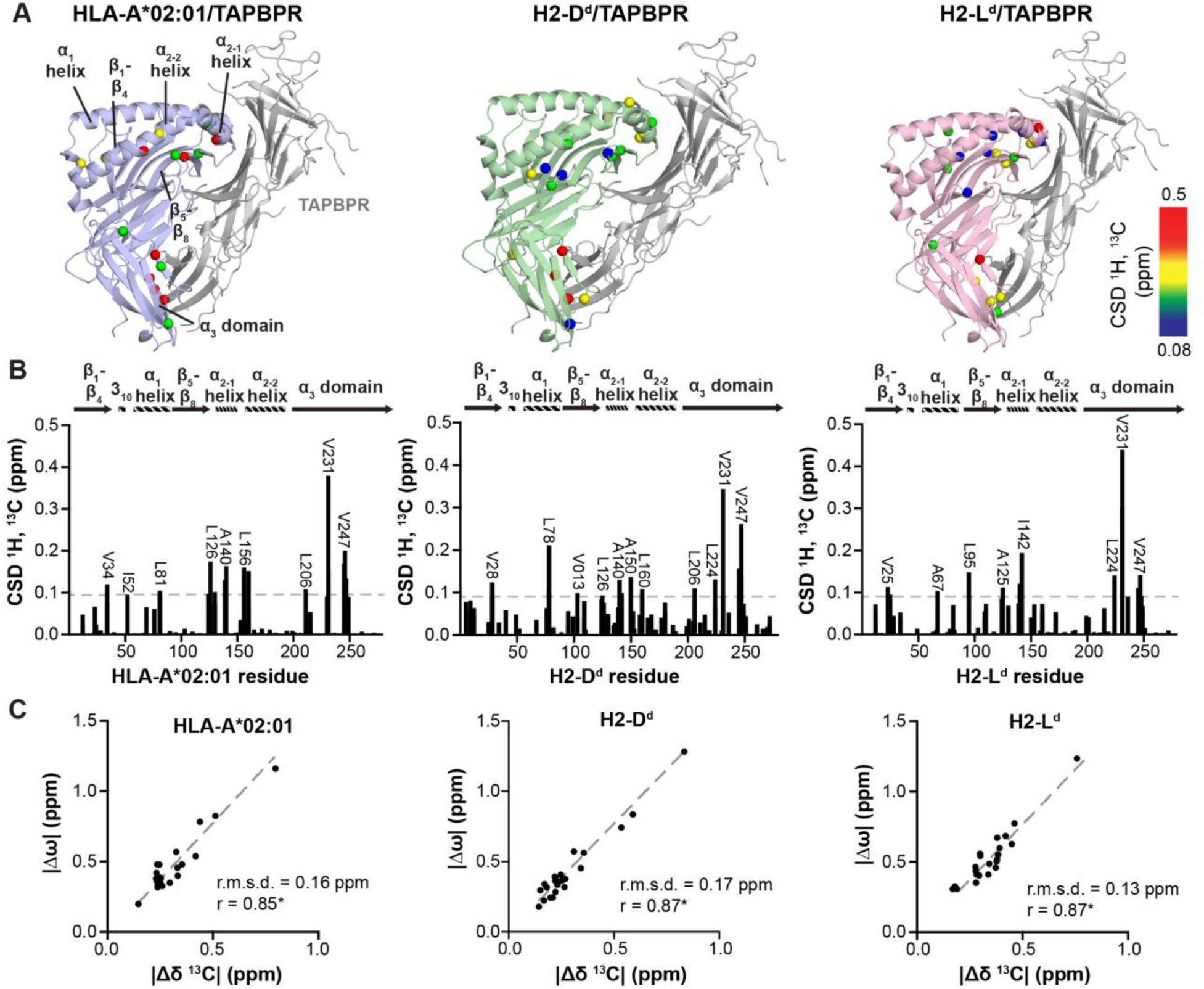
Dynamic surfaces of pMHC-I structures correlate with TAPBPR recognition sites. **(A)** Methyl chemical shift deviations (CSD) upon chaperone binding plotted onto the structure of MHC-I/TAPBPR complexes: TAX/HLA-A*02:01/hβ2m (blue), P18-I10/H2-D^d^/hβ2m (green) and NIH/H2-L^d^/hβ2m (pink) with TAPBPR shown in gray. The peptides are not shown. The H2-D^d^/TAPBPR crystal structure was obtained from PDB ID 5WER. The other structures are *Rosetta* homology models using PDB ID 5WER as a template (*85*). Affected regions are labeled. ΔCSDs are colored based on the scale shown at right. **(B)** Methyl CSDs measured from NMR titration experiments upon TAPBPR binding are shown as a function of heavy chain methyl residue number. Select methyl groups affected are noted. The gray dotted line represents the average CSD + 1 standard deviation. **(C)** The absolute value of the difference in the ^13^C chemical shift of the major and minor states of unchaperoned pMHC-I obtained from CPMG relaxation dispersion data (|Δω|, ppm) is shown as a function of the absolute value of the difference in the ^13^C chemical shift between the free and TAPBPR bound pMHC-I states determined from NMR titrations (|Δδ^13^C|, ppm). The slope (dotted gray line), the Pearson correlation coefficient (r), and the root mean square deviation (r.m.s.d.) are given for each correlation graph. The correlations are statistically significant with a P-value < 0.0001. HLA-A*01:01 is not shown because there is no detectable TAPBPR binding under the NMR sample conditions (see Fig. 2).

Our previous NMR study of the H2-D^d^ system suggested that MHC-I surfaces exhibiting μs-ms conformational dynamics in solution correlate with the TAPBPR recognition sites (*19*). To investigate whether this is a general trend, we compared heavy chain methyl groups displaying significant chemical shift changes upon complex formation (determined from NMR titrations) with those exhibiting μs-ms conformational exchange in the unchaperoned pMHC-I (determined from CPMG experiments) for the two additional interacting alleles, HLA-A*02:01 and H2-L^d^. A comparison of the absolute value of the difference between the ^13^C chemical shifts (|Δ^13^Cδ|) of the unchaperoned and TAPBPR-bound pMHC-I states versus |Δω| values, obtained from fits of CPMG dispersion profiles of free pMHC-I, uncovers a strong positive correlation (R^2^ 0.85-0.87) (**Fig. 4C**). Any conformational change in the pMHC-I that accompanies TAPBPR binding therefore occurs on pMHC-I surfaces of considerable structural plasticity in the free pMHC-I state.

Moreover, given that the |Δω| values directly report on conformational changes accompanying the formation of the minor (3-6% population) state sampled by free pMHC-I molecules in an aqueous solution, our observed quantitative correlation of |Δ^13^Cδ| with |Δω| supports a plausible model where the minor state is directly recognized by TAPBPR to form an initial encounter complex. Further structural adaptations, induced by direct interactions between the two proteins, then lead to the formation of the final, tight complex with each pMHC-I molecule. Since the extent of MHC-I regions participating in the formation of the minor state vary between MHC-I molecules of different heavy chain composition (**Fig. 3C-F),** with the non-interacting heavy chain HLA-A*01:01 exhibiting the most restricted dynamics, this model can explain allelic dependencies by differences in the minor state conformation.

### Restriction of the a_2-1_ helix abrogates sampling of the minor state and TAPBPR recognition

Our data, in conjunction with previous reports (*17*), provide strong evidence that the conformational plasticity of MHC-I molecules plays a key role in chaperone recognition. TAPBPR binding persists within the cell even in the presence of many disruptive mutations to the α_1_ domain, demonstrating that nascent MHC-I substrates for TAPBPR may be partially folded, with a high degree of conformational entropy (**Fig. S4A, B**). To further scrutinize this model in a controlled manner, we attempted to engineer an MHC-I molecule of restricted groove mobility. Such an engineered variant should also be stable for purification and detailed biophysical characterization in solution. We borrowed a design strategy previously applied to mouse H2-K^b^ to demonstrate that stabilization of the groove can rescue cell surface expression in a functional tapasin knockout (*57, 58*). A mutant MHC-I molecule, H2-D^d Y84C-A139C^, was prepared with an engineered disulfide between the sidechains of residues 84 and 139 across the F-pocket, which restricts the accessible conformational space for the MHC-I groove. Mouse H2-D^d^ was chosen as it is closely related to H2-K^b^, displays tight TAPBPR affinity in biophysical experiments, and finally because H2-D^d^ can still traffic to the plasma membrane when overexpressed together with suitable TAPBPR constructs. This property further allows us to distinguish between interactions with nascent intracellular MHC-I versus folded pMHC-I that escapes to the cell surface. By comparison, HLA-A*02:01 is retained internally (possibly in a nascent form) when co-expressed with TAPBPR (**Fig. S3A, B**). Soluble H2-D^d Y84C-A139C^, prepared as a complex with the P18-I10 peptide and human β2m, has comparable thermal stability to the wild-type molecule, suggesting that the disulfide does not compromise the pMHC-I structure (**Table S1**). Finally, we solved the crystal structure of the P18-I10/H2-D^d Y84C-A139C^/hβ2m complex at 2.37 Å resolution (**Table S3**, PDB ID 6NPR), which confirmed the presence of the oxidized C84-C139 disulfide bond and the bound P18-I10 peptide, and that the structure is very similar to the wild-type molecule (**Fig. 5A-C**).

**Fig. 5.**
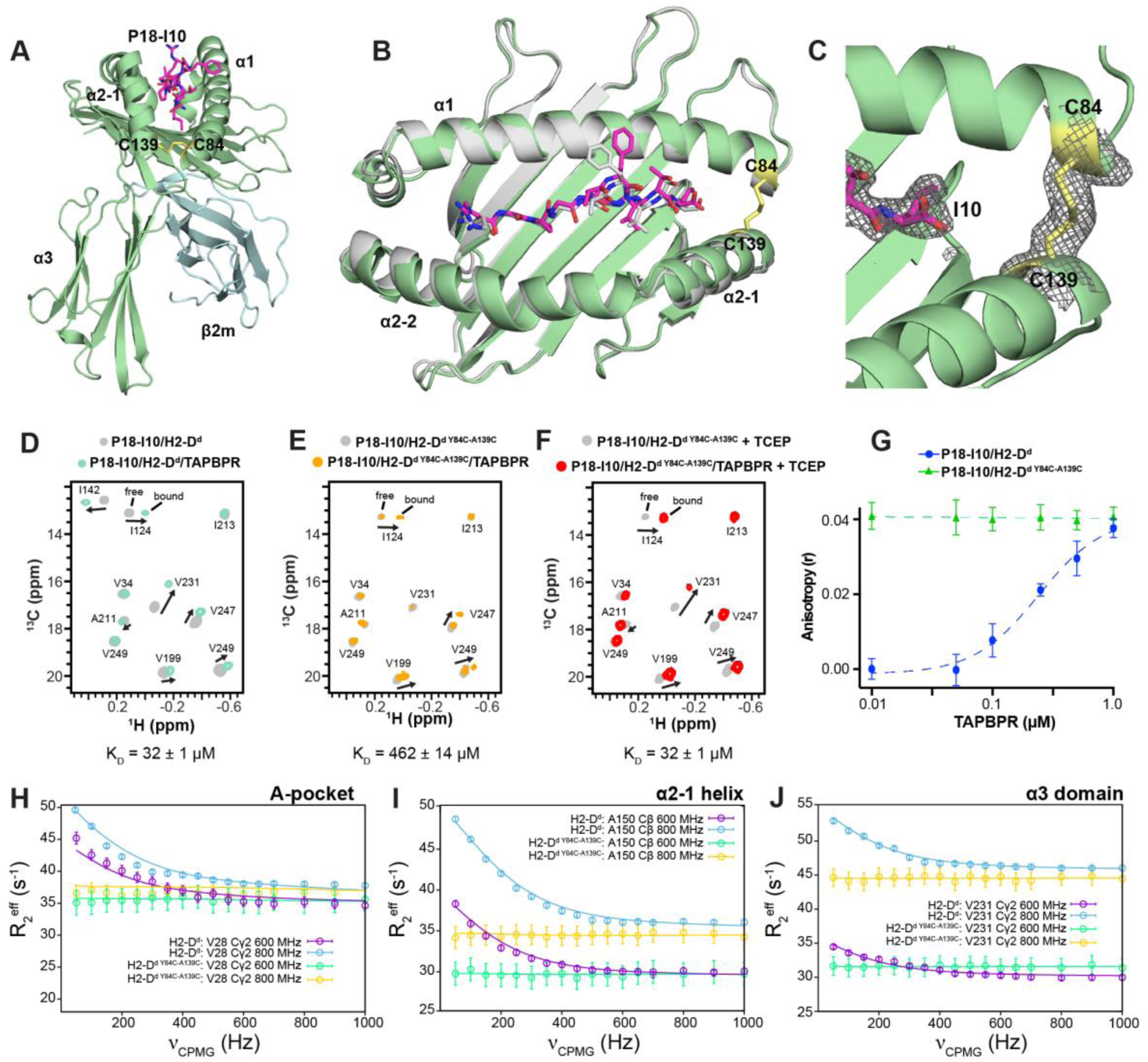
Restriction of dynamics in the pMHC-I groove abrogates binding to TAPBPR. **(A)** View of the P18-I10/H2-D^d Y84C-A139C^/hβ2m complex solved at 2.4 Å resolution (PDB ID 6NPR). The H2-D^d^ heavy chain is colored green, hβ2m cyan and P18-I10 magenta. The oxidized disulfide bond between C84 and C139 of H2-D^d^ is shown in yellow. **(B)** Overlay of the MHC-I groove (residues 1-180) and bound P18-I10 peptide for wild-type H2-D^d^ (PDB ID 3ECB, gray) and H2-D^d Y84C-A139C^ (colored as in A). The α_3_ domain and hβ2m are omitted for clarity. Backbone r.m.s.d. over the entire pMHC-I is 1.4 / 1.7 Å; (backbone / all-atom). **(C)** View of the P18-I10/H2-D^d Y84C-A139C^ structure showing the 2F_o_-F_c_ electron density map at 1.0 σ (gray mesh) around the P18-I10 peptide (magenta) and C84-C139 disulfide bond (yellow). **(D-F)** Representative 2D ^1^H-^13^C HMQC spectra from NMR titrations between TAPBPR and isotopically labeled (at the heavy chain) **(D)** wild-type P18-I10/H2-D^d^/hβ2m, **(E)** P18-I10/H2-D^d Y84C-A139C^/hβ2m, and **(F)** P18-I10/H2-D^d Y84C-A139C^/hβ2m in the presence of 1 mM TCEP. The NMR spectra shown were performed with three-fold molar excess TAPBPR. Dissociation constants obtained from NMR line shape fitting in TITAN are noted. **(G)** Fluorescence anisotropy experiments comparing exchange of TAMRA-labeled P18-I10 peptide with wild-type H2-D^d^ and H2-D^d Y84C-A139C^ as a function of TAPBPR concentration. **(H)**, **(I)**, **(J)** Comparison of representative ^13^C-SQ CPMG relaxation dispersion profiles for methyl groups of the heavy chain between wild-type P18-I10/H2-D^d^/hβ2m (800 MHz – blue; 600 MHz – purple) and P18-I10/H2-D^d Y84C-A139C^/hβ2m (800 MHz – yellow; 600 MHz – green) performed at 25°C. Both experiments were performed in the presence of three-fold molar excess P18-I10 peptide.

As predicted by our model, H2-D^d Y84C-A139C^ exhibits 15-fold lower affinity for TAPBPR relative to wild-type P18-I10/H2-D^d^/hβ2m by line shape analysis of NMR titrations performed using the program TITAN (**Fig. 5D, E**). The impaired interaction with TAPBPR is due to the presence of the C84-C139 disulfide, since specific reduction of this exposed disulfide with 1 mM of the mild reducing agent tris(2-carboxyethyl)phosphine (TCEP), further confirmed by NMR, rescues TAPBPR binding (**Fig. 5F**). As expected, the methyl chemical shifts of reduced H2-D^d Y84C-A139C^/TAPBPR complexes show no observable differences relative to the wild-type sample, suggesting that addition of small amounts of mild reducing agent does not affect the two conserved MHC-I disulfide bonds between C101-C164 and C203-C259, located at the buried interface of the α_2_ helix with the MHC platform and core of the α_3_ domain, respectively (**Fig. 5D-F**). In contrast to wild-type H2-D^d^, the disulfide-linked molecule demonstrates high levels of peptide exchange in fluorescence anisotropy experiments performed using a TAMRA-linked version of the P18-I10 peptide, even in the absence of TAPBPR (**Fig. 5G**), in agreement with recent reports that the same disulfide design approach can stabilize empty MHC-I molecules for a range of allelic compositions (*57, 58*). Notably, ^13^C-SQ CPMG relaxation dispersion experiments of P18-I10/H2-D^d Y84C-A139C^/hβ2m under the identical NMR conditions to experiments performed using wild-type H2-D^d^ clearly show that μs-ms conformational dynamics are quenched throughout the entire MHC-I structure, including the A-pocket, α_2-1_ helix and α_3_ domain regions, which exhibit clear dispersion profiles in the wild-type molecule **(Fig. 5H-J**). Together, these results further support our model of TAPBPR recognition through a network of dynamically coupled surfaces spanning multiple domains.

To complement these conclusions with experiments in the native cellular environment, tagged TAPBPR and H2-D^d^ constructs were co-expressed in human Expi293F cells. H2-D^d Y84C-A139C^ co-immunoprecipitated with TAPBPR similarly to wild-type H2-D^d^ (**Fig. S10**), despite showing reduced binding in the NMR experiments described above. Whereas binding interactions measured by NMR utilized purified, properly conformed H2-D^d Y84C-A139C^ in complex with the high-affinity P18-I10 peptide, within the cell nascent, empty MHC-I progressing through the secretory pathway are likely partially folded and exposing reduced cysteines. To resolve TAPBPR interactions with nascent/misfolded versus properly conformed MHC-I species, we compared wild-type TAPBPR to the TAPBPR-TM variant which can traffic to the plasma membrane, and measured interactions by BiFC. In agreement with the co-immunoprecipitation experiments, H2-D^d Y84C-A139C^ shows slightly enhanced BiFC with intracellular TAPBPR (**Fig. 6A, left**), presumably due to associations with nascent, partially reduced H2-D^d Y84C-A139C^ (**Fig. 6B**) (*5*). However, when folded H2-D^d Y84C-A139C^ molecules harboring a stabilizing disulfide are localized to the plasma membrane, association with TAPBPR-TM is substantially reduced, in agreement with our solution NMR results (**Fig. 6A, center**).

## Discussion

Our combined results suggest that TAPBPR has the capacity for multiple cellular functions: *(i)* a molecular chaperone of destabilized, empty MHC-I molecules at early stages of their folding process with broad substrate sequence specificity, and *(ii)* a peptide editor acting on properly conformed, peptide-loaded MHC-I molecules with a more exclusive specificity range (**Fig. 7**). The chaperone function may provide an important protein homeostasis mechanism, by stabilizing misfolded, aggregation-prone MHC-I species which form inside the cell (*5, 59*) and are known to contribute to the onset of disease (*60*). This cellular function is often attributed to a class of generic chaperones known as holdases, which includes essential heat-shock proteins (*61*). Our deep mutagenesis data suggest that the TAPBPR holdase activity requires a minimally folded epitope on the MHC α_2_ domain with a conserved disulfide (**Fig. 7A-C**), which is further consistent with the observed TAPBPR associations of a large panel of diverse MHC-I sequences and destabilizing mutants tested here. The role of the “editing” mode is likely to further scrutinize the peptide cargo of properly conformed MHC-I molecules during their egress to the cell surface. In this manner, TAPBPR helps to “focus” the antigen repertoire towards higher affinity peptides, such that pMHC-I molecules displayed at the cell surface exhibit sufficient stability and lifetime for immune recognition (*4, 5, 10, 11*). In contrast to holdase activity, the editing function of TAPBPR exhibits a narrow specificity range, where recognition of pMHC-I molecules is mediated by an excited state of 3 - 6% population in solution. This state is attained via a global conformational transition of the pMHC-I molecule as a whole through a network of allosterically coupled sites spanning both ends of the MHC groove and on the α_3_ domain. While the exact nature of the MHC-I conformations recognized by the chaperone and editing functions of TAPBPR remains to be fully described, our studies provide evidence that dynamics of the α_2_ domain are important for both recognition events. We propose that the chaperone and editing functions of TAPBPR have the same underlying structural basis. Based on crystal structures, TAPBPR binds to a conformation of the MHC-I α_2_ domain encompassing a widened groove. Nascent MHC-I molecules will readily sample multiple conformations, including those recognized by TAPBPR. However, as the MHC-I folds and binds to peptides, only MHC-I molecules that retain dynamic motions will sufficiently sample conformations selectively recognized by TAPBPR (**Fig. 7**). This property of the pMHC-I molecules is defined both by the allelic composition of the heavy chain, and the bound peptide.

Dynamic sampling of different MHC-I states has been previously suggested to be affected by both the bound peptide and the identity of amino acid sidechains in the D-, E- and F-pockets (*29, 30, 62*). Polymorphisms at specific positions, such as 114, 116, 147 and 156, have been shown to confer chaperone recognition of different alleles, due to these sites modulating the conformational flexibility and stability of peptide-deficient MHC-I (*21, 24, 62*). A sequence alignment of MHC-I molecules that recognize TAPBPR (H2-D^d^, H2-L^d^, HLA-A*02:01) versus one that does not (HLA-A*01:01) leads to no obvious clues to chaperone recognition, as the sites forming D-, E- and F-pockets for HLA-A*02:01 and HLA-A*01:01 are highly conserved (**Fig. S5**). Thus, future MD simulation experiments validated through measurements of dynamics by NMR and other biophysical techniques will be invaluable in understanding the exact molecular determinants that regulate MHC-I dynamics, given that protein dynamics involve transitions between multiple conformational basins with relatively low energy barriers, which can shift drastically in the presence of mutations due to allosteric effects (*63*).

A limitation of the present study is that the panel of heavy chain molecules tested has a very limited coverage of the more than 8,000 unique HLA sequences reported in the IPD-IMGT/HLA database (*64*). Additional experiments are needed to establish functionally important trends for TAPBPR, notably on the HLA-B and HLA-C classes. Equally important features of the MHC-I that contribute to the sequence-specificity of chaperone recognition remain to be characterized. First, with respect to the editing function of TAPBPR, bound peptides are known to contribute to the dynamic network on MHC-I structure (*19, 65, 66*). Second, the β2m light chain contributes to peptide binding, stabilization of the MHC-I groove, and forms direct contacts with TAPBPR (*14, 15, 19*). Although this is not addressed here, TAPBPR might also chaperone MHC-I molecules prior to their association with β2m, in a similar manner to tapasin (*67*). In our biophysical characterization of MHC-I/TAPBPR association, both human and mouse MHC-I molecules were prepared with human β2m. It remains unclear how heavy chain interactions with human versus mouse β2m, known to exhibit differences in affinity for the heavy chain, may modulate activity of the “editing” mode of TAPBPR (*68*). Furthermore, MHC-I molecules that are prepared in *E. coli* lack important glycan modifications present *in vivo*, which could further contribute to chaperone recognition, either via direct interactions or due to changes in the dynamic landscape of the MHC-I (*64, 65*). Finally, we have interpreted our cellular data on the basis of direct interactions between TAPBPR and MHC-I, due to their known physical association and because the HLA-A*02:01 mutational landscape is consistent with crystal structures. However, we cannot yet definitively rule out that allele specificity is broadened within the cell due to indirect associations mediated by higher-order chaperone assemblies.

In summary, our data provide insights into the conformational plasticity of different MHC-I molecules by a range of techniques, and suggest a plausible role of these coupled motions in shaping the allelic specificity of TAPBPR recognition. The conserved MHC-I recognition mechanism shared between TAPBPR and tapasin implies that a similar paradigm of dynamically driven recognition may also influence MHC-I allelic dependencies for tapasin in the peptide-loading complex. These findings are a first step in elucidating how molecular chaperones selectively edit the displayed antigen repertoire for different alleles, towards understanding their role in the highly complex associations between HLA polymorphisms and human diseases.

## Materials and Methods

Specific details about protein expression and purification, ITC experiments NMR assignments, chemical shift mapping and CPMG relaxation dispersion experiments, X-ray crystallography, fluorescence anisotropy, co-immunoprecipitation/immunoblots and bimolecular fluorescence complementation/deep sequencing assays are outlined in detail in the Supporting Information.

## Data and Materials Availability

Raw and processed Illumina sequencing data are deposited with NCBI’s Gene Expression Omnibus under series accession number GSE128957. Reviewers can access the private data using token: cdktmweknpspvsp. NMR assignments have been deposited into the Biological Magnetic Resonance Data Bank (http://www.bmrb.wisc.edu) under accession numbers 27249 [H2-D^d^], 27682 [H2-L^d^], 27632 [HLA-A*01:01] and 27631 [HLA-A*02:01]. The atomic coordinates and structure factors for the P18-I10/H2-D^d Y84C-A139C^/β2m complex have been deposited in the Protein Data Bank (https://www.rcsb.org/) under the PDB ID 6NPR. The other pMHC-I structures used in this study were published previously: NRAS^Q61K^/HLA-A*01:01 (PDB ID 6MPP); TAX/HLA-A*02:01 (PDB ID 1DUZ) (*44*); P18-I10/H2-D^d^ (PDB ID 3ECB) (*42*); NIH**/**H2-L^d^ (PDB ID 1LD9) (*41*); H2-D^d^/TAPBPR (PDB IDB 5WER). Plasmids will be available from Addgene (https://www.addgene.org) upon manuscript publication. Further information and requests for resources and reagents should be directed to and will be fulfilled by the Lead Contact, Nikolaos G. Sgourakis.

## Author Contributions

N.G.S. conceived, and together with E.P. designed the project. A.C.M. produced labelled MHC-I samples, performed SEC, NMR, ITC and DSF experiments, Rosetta modeling and data analysis. C.A.D., J.P., and E.P. produced MHC-I sequence libraries, performed confocal microscopy, co-immunoprecipitation, bimolecular fluorescence complementation experiments and data analysis. H.C. performed immunoblots. S.A.O. and D.M. produced recombinant TAPBPR samples. J.S.T. performed fluorescence anisotropy experiments and together with S.T. solved the X-ray structure of the disulfide H2-D^d^ mutant. A.C.M., S.A.O. and D.F.S. produced NMR assignments. A.C.M., E.P. and N.G.S. wrote the manuscript, with feedback from all authors.

## Acknowledgements

This research was supported by NIAID grant (1R01AI129719) to E.P., by NIAID (AI2573) and NIGMS (1R35GM125034) grants to N.G.S., and by a High End Instrumentation (HIE) Grant (S10OD018455) which funded the 800 MHz NMR spectrometer at UCSC. The authors acknowledge the resources of the Advanced Photon Source, beamline 23-ID-B, a U.S. Department of Energy (DOE) Office of Science User Facility operated for the DOE Office of Science by Argonne National Laboratory under Contract No. DE-AC02-06CH11357. We acknowledge the Lewis Kay (University of Toronto) lab for providing the SQ ^13^C-CPMG pulse sequences, Flemming Hansen (University College London) for aid with CPMG fitting in CATIA, Hsiau-Wei Lee (UCSC) for assistance in acquiring NMR experiments at 600 MHz, Kannan Natarajan (NIAID) for providing the S2 cell lines for TAPBPR protein expression and Tilini Wijeratne and Giora Morozov (UCSC) for help with protein expression and purification. Cell sorting and deep sequencing were conducted at the Roy J. Carver Biotechnology Center (Illinois) with assistance from Barbara Pilas, Barbara Balhan, Christy Wright, and Alvaro Hernandez. We are thankful to Kannan Natarajan and David Margulies (NIAID) for helpful discussions throughout the course of this study.

## Supporting Information

### Supporting Methods

#### Expression Constructs for Cell Assays

Expi293F cells (ThermoFisher) were cultured in Expi293 Expression Medium (ThermoFisher) at 37 °C, 125 rpm, and 8% CO_2_. FLAG-TAPBPR (containing from N- to C-terminus the signal peptide of influenza HA, a FLAG tag, a GGS linker, and TAPBPR residues K1-S447) was PCR assembled. TAPBPR-TM was assembled by fusing the ectodomain of FLAG-TAPBPR up to residue R384, followed by a SGAGSA linker, on to amino acids (a.a.) I282-D314 of HLA-G encoding a canonical transmembrane domain. Tagged MHC-I alleles were constructed featuring from N- to C-terminus the signal peptide of influenza HA, a c-myc tag, a GSPGGSSGGG linker, and mature MHC-I. H2-D^d^ constructs carried the C121S mutation, removing a non-oxidized extracellular cysteine residue to prevent spurious disulfide formation during engineering. For BiFC experiments, the C-termini of FLAG-TAPBPR and FLAG-TAPBPR-TM were fused to VC (a.a. D155–K238 of Venus). The C-termini of myc-tagged MHC-I were fused to VN (a.a. V1–A154 of Venus I152L mutant) (*71*). Cloning of tagged CXCR4 fusions to VN and VC are previously described (*72*). All constructs were cloned into pCEP4 (Invitrogen), and targeted mutations were generated by site directed mutagenesis and confirmed with DNA sequencing. Plasmids will be available from Addgene upon publication.

#### Bimolecular Fluorescence Complementation

Expi293F cells at 2×10^6^ per mL were co-transfected with 500 ng of plasmids encoding the respective VN- and VC-fusions using ExpiFectamine (ThermoFisher). Cells were harvested 23-25 h post transfection and fixed using BD Fixation/Permeabilization kit (BD Biosciences). Cells were washed with BD Perm/Wash Buffer, stained with anti-FLAG Cy3 (clone M2, 1/200 dilution; Sigma-Aldrich) and anti-myc Alexa 647 (clone 9B11, 1/200 dilution; Cell Signaling Technology), washed twice, and resuspended in Dulbecco’s phosphate-buffered saline (PBS) supplemented with 0.2% bovine serum albumin (BSA) for analysis on a BD LSR II with three-color compensation. Data were analyzed with FCS Express 6 (De Novo Software).

#### Immunoblots

Cell samples were denatured in reducing sodium dodecyl sulfate (SDS) load dye, and proteins were separated by gel electrophoresis and transferred to polyvinylidene difluoride membrane. For FLAG-tagged proteins, membranes were blocked with 3% BSA in Tris-buffered saline-0.1% Tween 20 (TBST), incubated in 1/2,000 anti-FLAG (M2) AP (Sigma-Aldrich), washed thoroughly and detected with 1-Step NBT-BCIP (ThermoFisher). Blots for myc-tagged proteins were blocked with 5% BSA in TBST, incubated in 1/2,000 anti-myc (71D10) HRP (Cell Signaling Technology), and detected using Clarity Western ECL substrate (Bio-Rad).

#### Immunoprecipitations

Transfected Expi293F cells were harvested 23-25 h post transfection and freeze-thawed twice in PBS supplemented with protease inhibitors (Sigma-Aldrich). Lysed samples were centrifuged at 21,000 g. Membrane pellets were solubilized with precipitation buffer (50 mM Tris pH 7.5, 150 mM NaCl, 5 mM MgCl_2_, 1 mM EDTA) containing 0.5% dodecylmaltoside (DDM; Anatrace). Insoluble debris was removed by centrifugation at 21,000 g. The soluble fraction was incubated with FLAG M2 resin (Sigma-Aldrich) for 1 h at 4 °C, and then washed three times with precipitation buffer containing 0.05% DDM. Resin was resuspended in reducing SDS load dye for immunoblotting.

#### Library Generation

Using the pCEP4-myc-HLA-A*02:01-VN expression plasmid, a SSM library focused on the α_1_/α_2_ domains (a.a. S2-R181) was created using overlap extension PCR (*73*). The library covered 3,524 of 3,600 possible single amino acid substitutions based on a minimum frequency of 2 × 10^−6^. Expi293F cells at 2 × 10^6^ cells/mL were transfected using Expifectamine (ThermoFisher) with 1 ng library DNA diluted with 1.5 μg of pCEP4-ΔCMV (*74*). The media was replaced 2 hours post transfection. Under these conditions, cells typically acquire no more than a single coding sequence (*72*).

#### Sorting for Myc-HLA-A*02:01 Surface Expression

Expi293F cells were harvested 25 h post-transfection. Cells were washed on ice with PBS-BSA, stained with Alexa Fluor 647-conjugated anti-myc clone 9B11 (1:200 dilution; Cell Signaling Technology), washed twice and resuspended in PBS-BSA. Cells were stained with propidium iodide (final concentration of 1 μg/mL) immediately prior to sorting. Cells were gated by scattering properties to exclude debris and doublets, and propidium iodide-positive cells were also excluded. The top 0.5% of cells for Alexa Fluor 647 fluorescence were collected into fetal bovine serum coated tubes containing media during a 4 h sort on a BD FACSAria II. Collected cells were pelleted and frozen at −80 °C. Replicate selections were performed independently on different days.

#### Sorting for BiFC Signal

Expi293F cells were transfected with pCEP4-FLAG-TAPBPR-TM-VC linearized by EcoRV, selected with 100 μg/ml hygromycin B, and FACS sorted for FLAG positive cells. The resulting stable line was transfected with the library as described above. Cells were harvested 25 h post-transfection, washed and resuspended in ice-cold PBS-BSA. Cells were sorted using a BD FACSAria II, excluding debris and doublets based on scattering properties. The top 20% of Venus-positive cells (equivalent to the top 1% of the total population) were collected and frozen at −80 °C.

#### Deep Sequencing

RNA was extracted from sorted cells using the GeneJet RNA purification kit (Fisher). cDNA was prepared using Accuscript reverse transcriptase (Agilent) with a primer that annealed to the VN region to prevent amplification of endogenous HLA-A*02:01 transcripts. DNA fragments spanning the mutated region were PCR amplified in two rounds: the first appended sequences complementary to Illumina sequencing primers, and the second added experiment-specific barcodes and Illumina adaptamers. Sequencing was on a NovaSeq 6000 and analyzed with Enrich (*75*). Data deposited with NCBI’s Gene Expression Omnibus under series accession number GSE128957 includes primer sequences and commands. Reviewers can access the private data using token: cdktmweknpspvsp

#### Confocal Microscopy

Performed as described in (*74*). Briefly, transfected HEK293T cells were permeabilized and fixed using the BD Cytofix/Cytoperm kit, and co-stained with anti-FLAG-Cy3 (Sigma-Aldrich) and anti-GM130-Alexa Fluor 488 (ThermoFisher Scientific) or anti-calnexin-Alexa Fluor 488 (ThermoFisher Scientific). Images were collected on a Zeiss LSM 700 (Carl Zeiss) and processed using Fiji.

#### Expression Constructs for Protein Purification

DNA encoding the luminal domain of MHC-I heavy chains of murine H2-D^d^ and H2-L^d^, and human HLA-A*02:01 and HLA-A*01:01, were transformed into *Escherichia coli* BL21(*DE3)* (Novagen). MHC-I molecules were expressed in Luria-Broth, extracted from inclusion bodies, refolded *in vitro* at 4°C and pMHC-I complexes were purified as previously described (*19*). Known full-length peptide antigens used for refolding were prepared by chemical synthesis (Biopeptek Inc, Malvern, USA or GenScript, Piscataway, USA): NIH peptide (YPNVNIHNF) for H2-L^d^ (*41*); P18-I10 peptide (RGPGRAFVTI) for H2-D^d^ (*42*); NRAS^Q61K^ peptide (ILDTAGKEEY) for HLA-A*01:01 (*43*); TAX peptide (LLFGYPVYV) for HLA-A*02:01 (*44*). The luminal domain of TAPBPR was expressed using a Drosophila S2 cell expression system and purified as previously described (*13*). All purified proteins were exhaustively buffer exchanged into 100 mM NaCl, 20 mM sodium phosphate pH 7.2.

#### X-ray crystallography

P18-I10/H2-D^d Y84C-A139C^/hβ2m crystals were grown by the sitting drop method and plates were incubated at 22°C. Crystals were obtained by mixing 1 μL of protein at 12 mg/mL with an equal volume of reservoir solution (0.2M ammonium acetate, 0.1M Bis-Tris pH 5.5 and 25% (v/v) PEG 3350). Reservoir solution supplemented with 20% (v/v) glycerol used as a cryoprotectant. Data were collected at the Advanced Photon Source, Argonne National Laboratory, at beamline 23-IDB. Diffraction images were indexed, integrated, and scaled using Mosflm and Aimless in the CCP4 package (*76*). Structures were determined by Phaser (*77*) using PDB ID 3ECB as a search model. Model building and refinement were performed using COOT (*78*) and Phenix (*79*), respectively. The structural model and structure factors were deposited into the Protein Data Bank under accession code 6NPR.

#### Differential Scanning Fluorimetry

DSF experiments were performed on an Applied Biosystems ViiA 7 qPCR machine with excitation and emission wavelengths set to 470 nm and 569 nm with proteins in buffer of 100 mM NaCl, 20 mM sodium phosphate pH 7.2. Experiments were conducted in triplicate in MicroAmp Fast 96-well plates with 50 μL total volume containing final concentrations of 7 μM protein and 10× SYPRO orange dye (ThermoFisher). Temperature was incrementally increased at a scan rate of 1°C/min between 25°C and 95°C. Data analysis and fitting were performed in GraphPad Prism v7.

#### SEC Binding Assays

Size exclusion chromatography (SEC) binding assays were performed using a mixture of 80 µM pMHC-I and 80 µM TAPBPR (1:1 molar ratio) that were incubated at room temperature for 1 hour. SEC binding was performed on a Superdex 200 Increase 10/300 GL column at flow rate 0.5 mL/min in 100 mM NaCl, 20 mM sodium phosphate pH 7.2.

#### ITC experiments

Isothermal titration calorimetry (ITC) experiments between pMHC-I and TAPBPR constructs were obtained using a MicroCal VP-ITC system (Malvern Panalytical). All proteins were exhaustively dialyzed into the buffer (50 mM NaCl, 20 mM sodium phosphate pH 7.2) filtered through a 0.22 μm PES membrane. Syringe containing pMHC-I at concentrations of 100 to 150 μM were titrated into calorimetry cell containing 12 μM TAPBPR and 1 mM purified free peptide (NRAS^Q61K^ for HLA-A*01:01, TAX for HLA-A*02:01, P18-I10 for H2-D^d^ and p29 for H2-L^d^). Injection volumes were 10 μL performed for a duration of 10 sec and spaced 220 sec apart to allow for a complete return to baseline. Data was subtracted from a control performed where syringe containing pMHC-I at concentrations of 100 to 150 μM were titrated into calorimetry cell containing buffer and 1 mM purified free peptide. Data were processed and analyzed with Origin software. Isotherms were fit using a one-site ITC binding model. The first data point was excluded from analysis. Reported K_D_, -T*ΔS, and ΔH values are the average values from two technical replicates.

#### NMR Chemical Shift Assignments

Samples for NMR were prepared using AILV methyl (Ala ^13^Cβ, Ile ^13^Cδ1, Leu ^13^Cδ1/^13^Cδ2, Val ^13^Cγ1/^13^Cγ2) labeling at the heavy chain of H2-D^d^, H2-L^d^, HLA-A*02:01 or HLA-A*01:01 pMHC-I molecules, on a ^12^C/^2^H/^15^N background (*19*). Resonance assignments were derived separately for each pMHC-I system from a series of 3D experiments recorded at 800 MHz and 25°C (*19, 47*). Briefly, backbone amide resonances were assigned using standard, TROSY-based 3D HNCO, HNCA and HN(CA)CB experiments recorded at 800 MHz at 25°C. Backbone amide assignments were further validated through amide-amide NOEs, acquired in 3D H_N_-NH_N_ and 3D N-NH_N_ SOFAST NOESY experiments (3). Unambiguous assignments of methyl resonances were obtained on the basis of the backbone assignments, using 3D HMCM[CG]CBCA out-and-back methyl-TROSY experiments recorded on 500 μM to 1 mM pMHC-I samples prepared with an alternative ILV methyl labelling scheme (ILV*) which aims to generate a linear ^13^C labelling pattern at the side chains of Leu and Val spin systems for optimal sensitivity (*81*). AILV methyl assignments were validated and stereo-specifically disambiguated by acquiring methyl-methyl NOEs in 3D H_M_-C_M_H_M_ and 3D C_M_-C_M_H_M_ SOFAST NOESY experiments (*80*). All 3D SOFAST NOESY experiments were acquired using standard parameters (*47*). To confirm the assignment of the TAPBPR bound pMHC-I states for the sidechain methyl resonances, an additional 3D H_M_-C_M_H_M_ experiment was acquired with 200 µM pMHC-I/TAPBPR complexes and compared to similar NOE strips from the unbound pMHC-I reference. All NMR data were processed with NMRPipe (4) and analyzed using NMRFAM-SPARKY (5).

#### NMR Titrations and Chemical Shift Mapping

NMR chemical shift mapping of different pMHC-I/TAPBPR complexes was performed in a manner analogous to our established characterization of the H2-D^d^ system (*19*). Briefly, 105 µM pMHC-I was titrated with increasing concentrations of unlabeled TAPBPR in matched NMR buffer (100 mM NaCl, 20 mM sodium phosphate pH 7.2, 10% D_2_O (v/v), 1U Roche protease inhibitor). pMHC-I:TAPBPR molar ratios were 1:0, 1:0.59, 1:1.18, 1:1.77, 1:2.36, and 1:2.95. Two-dimensional ^1^H-^13^C methyl SOFAST HMQC experiments recorded at 25°C at a ^1^H field strength of 800 MHz were used as readouts. A total number of 136 scans were used with a 0.2 sec recycle delay (d1) and acquisition times of 30 ms in the ^13^C dimension. To compare with the SOFAST experiments, an additional titration between 105 µM pMHC-I with pMHC-I:TAPBPR molar ratios were 1:0, 1:0.59, 1:1.18, 1:1.77, 1:2.36, and 1:2.95 was performed using standard (non-SOFAST) version of the 2D ^1^H-^13^C HMQC experiment recorded at 25°C at a ^1^H field strength of 800 MHz with 28 scans were used with a 0.8 sec recycle delay (d1) and acquisition times of 30 ms in the ^13^C dimension. Data were processed with 4 Hz and 10 Hz Lorentzian line broadening in the direct and indirect dimensions, respectively, and fit using a two-state binding model in TITAN (*84*) with bootstrap error analysis of 100 replicas. Chemical shift deviations (CSD, p.p.m.) were determined using the equation Δδ^CH3^=[1/2(Δδ_H_^2^+Δδ_C_^2^/4)]^1/2^ for ^13^C AILV methyls. The |Δδ ^13^C| was determined by taking the absolute value of the difference between the ^13^C chemical shift of the free and TAPBPR bound pMHC-I states.

#### CPMG Relaxation Dispersion

AILV methyl sidechain relaxation dispersion profiles were obtained using ^1^H to ^13^C single-quantum Carr-Purcell-Meiboom-Gill (CPMG) relaxation dispersion experiments (*53*) on samples with concentrations ranging from 500 μM to 1 mM perdeuterated pMHC-I samples recorded at 25°C at 600 MHz and 800 MHz. A temperature calibration using the temperature-dependence of the water resonance, relative to an internal 4,4-demethyl-4-silapentane-1-sulfonic acid (DSS) standard was used to ensure strict temperature matching between the two instruments operating at different magnetic fields. Spectra were recorded in an interleaved manner with CPMG field strengths of 50, 100, 150, 200, 250, 300, 350, 400, 450, 500, 550, 600, 650, 700, 800, 900 and 1000 Hz with a constant time delay (T_CPMG_) of 40 ms. Peak intensities obtained using NMRFAM-SPARKY were converted to the R_2, eff_ transverse decay rates with the equation R_2, eff_ = 1/T_CPMG_ × ln (I_0_ / I_νCPMG_). Only with non-overlapping resonances were analyzed. CPMG profiles were fitted with a two-state, global model of all methyl groups displaying dispersion (R_ex_ > 1 s^−1^) using the program CATIA, which further allows for a correction of off-resonance effects of the CPMG 180^°^ pulse train (http://www.biochem.ucl.ac.uk/hansen/catia/).

#### Fluorescence Anisotropy

Fluorescence anisotropy (r, herein referred to as FA) was performed using a P18-I10 peptide labeled with TAMRA dye (K^TAMRA^GPGRAFVTI, herein called TAMRA-P18-I10) (Biopeptek Inc.) (*19*). Graded concentrations (0 μM, 0.05 μM, 0.1 μM, 0.25 μM, 0.5 μM and 1 μM) of TAPBPR were added to a mixture of 0.75 nM TAMRA-P18-I10 and either 0.1 μM of wild-type P18-I10/H2-D^d^/hβ2m or P18-I10/H2-D^d Y84C-A139C^/hβ2m. The average FA after incubation for 95–105 min at room temperature (25 °C) was plotted as a function of the log_10_ of concentration of TAPBPR. Each experiment was performed at room temperature in a volume of 100 μL and loaded onto a black 96-well polystyrene assay plate (Costar 3915). FA data were recorded via a PerkinElmer Envision 2103 plate reader with an excitation filter of λ_ex_ = 531 nm and an emission filter of λ_em_ = 595 nm. Each experiment was performed in triplicate. Experimental values were subtracted from background FA values obtained from incubation of TAMRA-P18-I10 alone. All samples were prepared in matched buffer (100 mM NaCl, 20 mM sodium phosphate pH 7.2, 0.05% (v/v) tween-20). Data analysis and fitting was performed in GraphPad Prism v7.

#### Rosetta modeling

*Rosetta*’s comparative modeling protocol was used to generate MHC-I/TAPBPR complexes using the H2-D^d^/β2m/TAPBPR crystal structure as a template (*15, 85*). High resolution structure refinement of the resulting models were carried out using *Rosetta*’s relax protocol (*86*).

## Supporting Figures

**Fig. S1.**
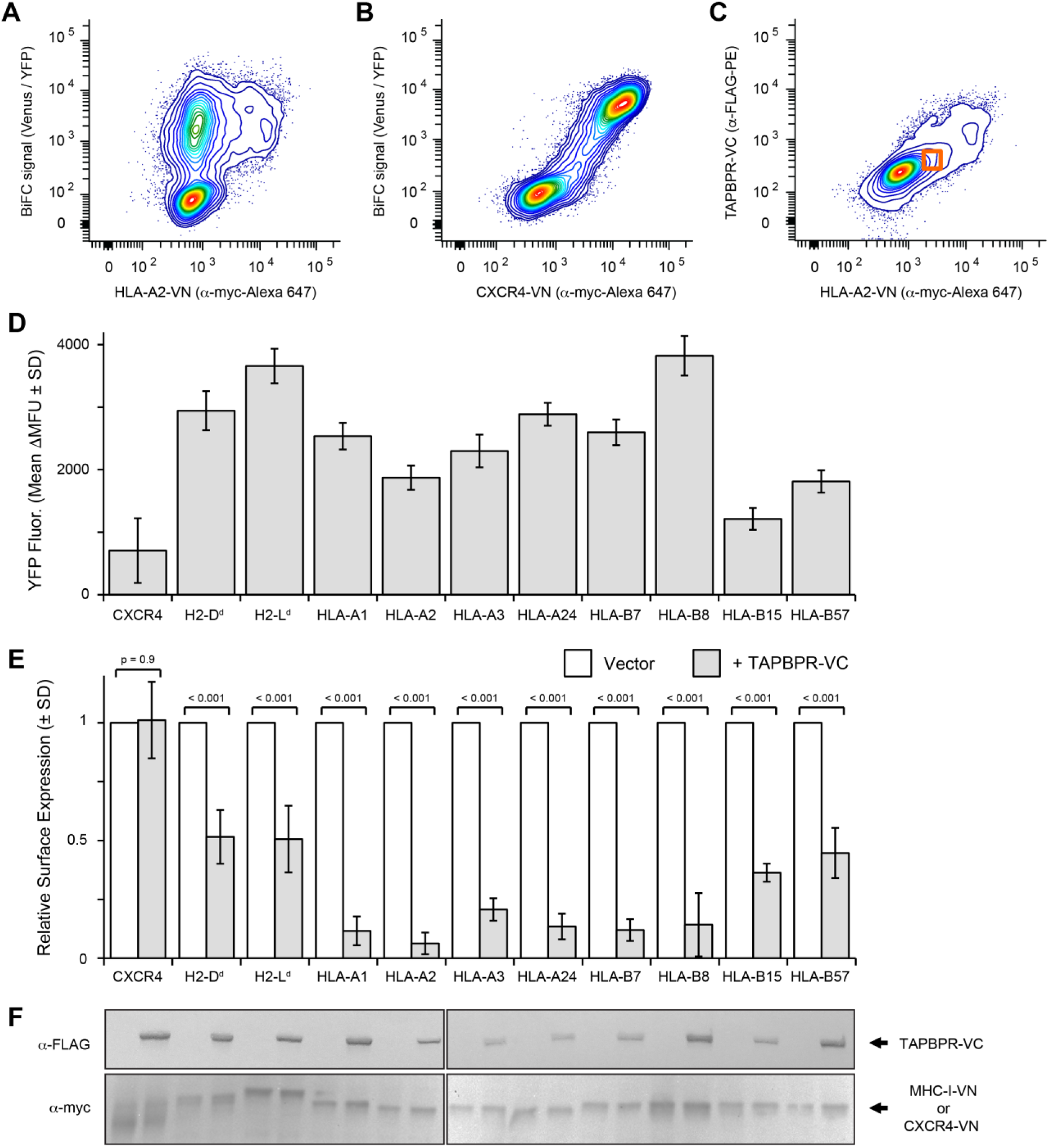
TAPBPR associates with a broad range of MHC-I alleles in a native cellular environment. **(A)** Intracellular staining and flow cytometry analysis of Expi293F cells co-expressing myc-HLA-A*02:01-VN (*x*-axis) and FLAG-TAPBPR-VC. Venus fluorescence (*y*-axis) is high at low HLA-A*02:01 expression. **(B)** FLAG-TAPBPR-VC was co-expressed with myc-CXCR4-VN (*x*-axis) as a negative control. Venus fluorescence (*y*-axis) is only elevated at very high CXCR4 expression due to non-specific interactions. **(C)** To partially compensate for expression differences and exclude highly expressing cells with saturated BiFC, cells were gated (orange box) for low MHC-I or CXCR4 (*x*-axis) and TAPBPR (*y*-axis) expression prior to measuring BiFC signal. **(D)** BiFC between TAPBPR-VC and CXCR4-VN or MHC-I-VN. Average ΔMean Fluorescence Units (ΔMFU) ± SD, n = 4. **(E)** Surface expression levels of myc-tagged VN constructs in the absence and presence of TAPBPR-VC were determined by flow cytometry. Mean ± SD, n = 7, *p* values determined by two-tailed Student’s t test. In panels **(D)** and **(E)** the full allele names for HLA molecules tested are HLA-A*01:01, HLA-A*02:01, HLA-A*03:01, HLA-A*24:02, HLA-B*07:02, HLA-B*08:01, HLA-B*15:01 and HLA-B57:01. **(F)** Western blots for a representative experiment comparing total protein expression levels for TAPBPR (α-FLAG) and CXCR4 or heavy chain MHC-I (α-myc) constructs. Samples are vertically aligned with graphs above.

**Fig. S2.**
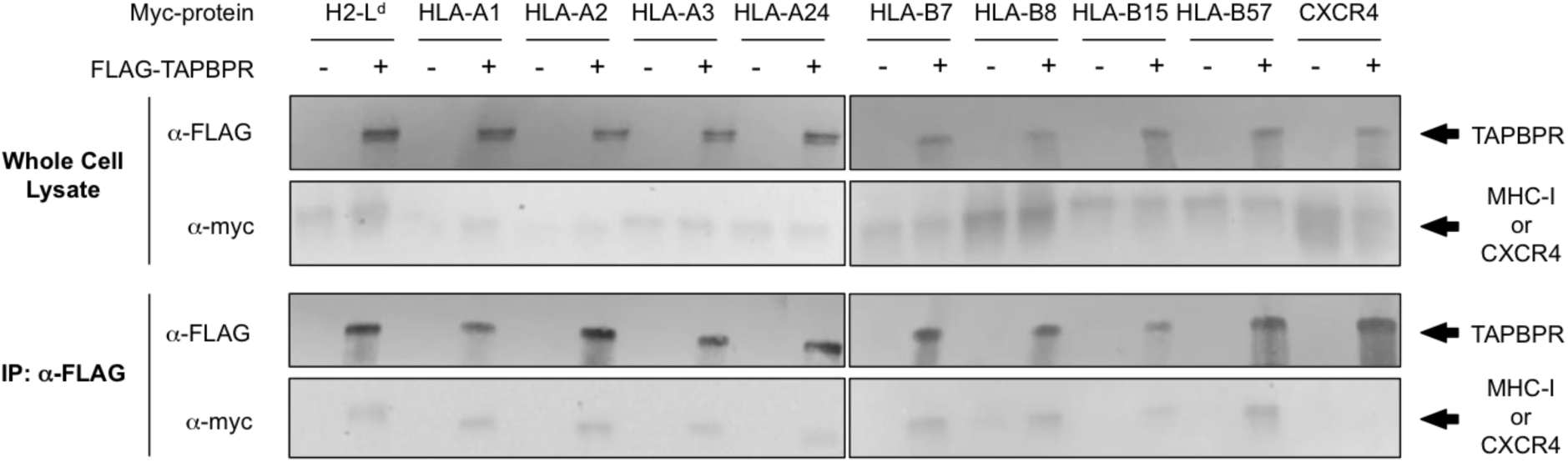
TAPBPR co-immunoprecipitates with a broad range of MHC-I alleles. FLAG-TAPBPR was co-expressed with myc-MHC-I or the negative control myc-CXCR4 (see upper immunoblots of protein in whole cell lysates). TAPBPR was precipitated with anti-FLAG resin and bound proteins were detected (lower immunoblots). Shown are representative results from two replicates. The full allele names for HLA molecules tested are HLA-A*01:01, HLA-A*02:01, HLA-A*03:01, HLA-A*24:02, HLA-B*07:02, HLA-B*08:01, HLA-B*15:01 and HLA-B57:01. Refer to Fig. S10 for immunoprecipitations of H2-D^d^ with TAPBPR.

**Fig. S3.**
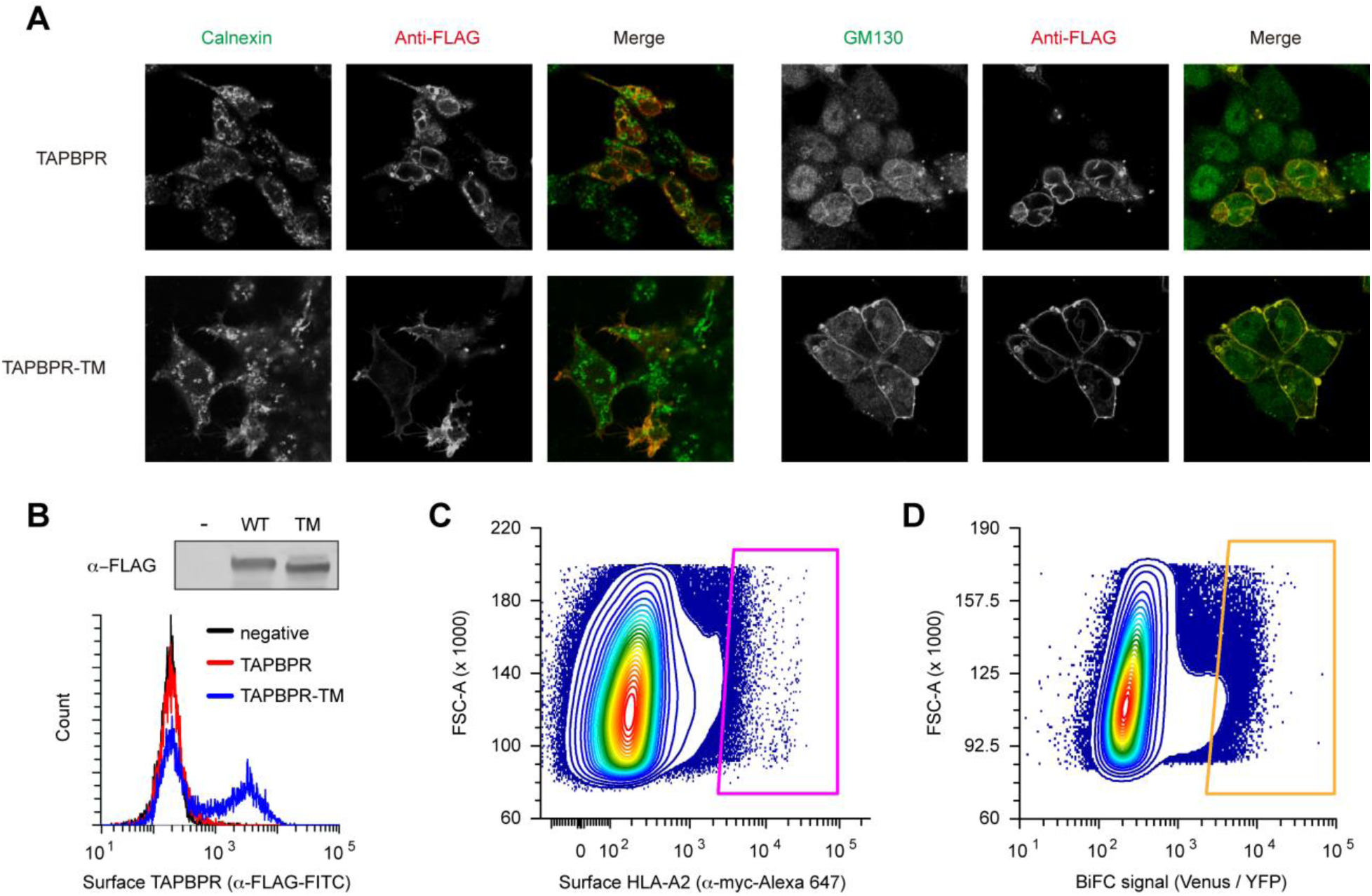
Replacement of the TAPBPR transmembrane domain facilitates escape to the cell surface. **(A)** Confocal microscopy of transfected HEK293 cells shows that both wild-type TAPBPR and intracellular TAPBPR-TM colocalize with Golgi marker GM130, but less so with the ER protein calnexin. The TAPBPR constructs are FLAG tagged at their extracellular/luminal N-termini. **(B)** Flow cytometry analysis of surface expressed wild-type TAPBPR (red) or TAPBPR-TM (blue) in transfected Expi293F cells. Both proteins are detected by immunoblot of whole lysate (upper inset). **(C)** For selection of HLA-A*02:01 sequence variants that are surface expressed, a saturation mutagenesis library focused on the α_1_/α_2_ domains was constructed on HLA-A*02:01 featuring an N-terminal myc tag for surface detection and a C-terminal VN fusion for BiFC. The plasmid library was transfected in to Expi293F cells such that typically no more than a single variant is acquired by any cell. Under these conditions, most cells remain negative. Cells highly expressing surface-localized HLA-A*02:01 were collected by FACS (shown by the magenta gate, top 0.5 % of population). **(D)** For selection of HLA-A*02:01 variants competent for TAPBPR interactions, the HLA-A*02:01 library was now transfected in to Expi293F cells stably expressing TAPBPR-TM fused to VC. The top 1% of total cells for BiFC signal (orange gate) were collected, equivalent to 20 % of the Venus-positive population.

**Fig. S4.**
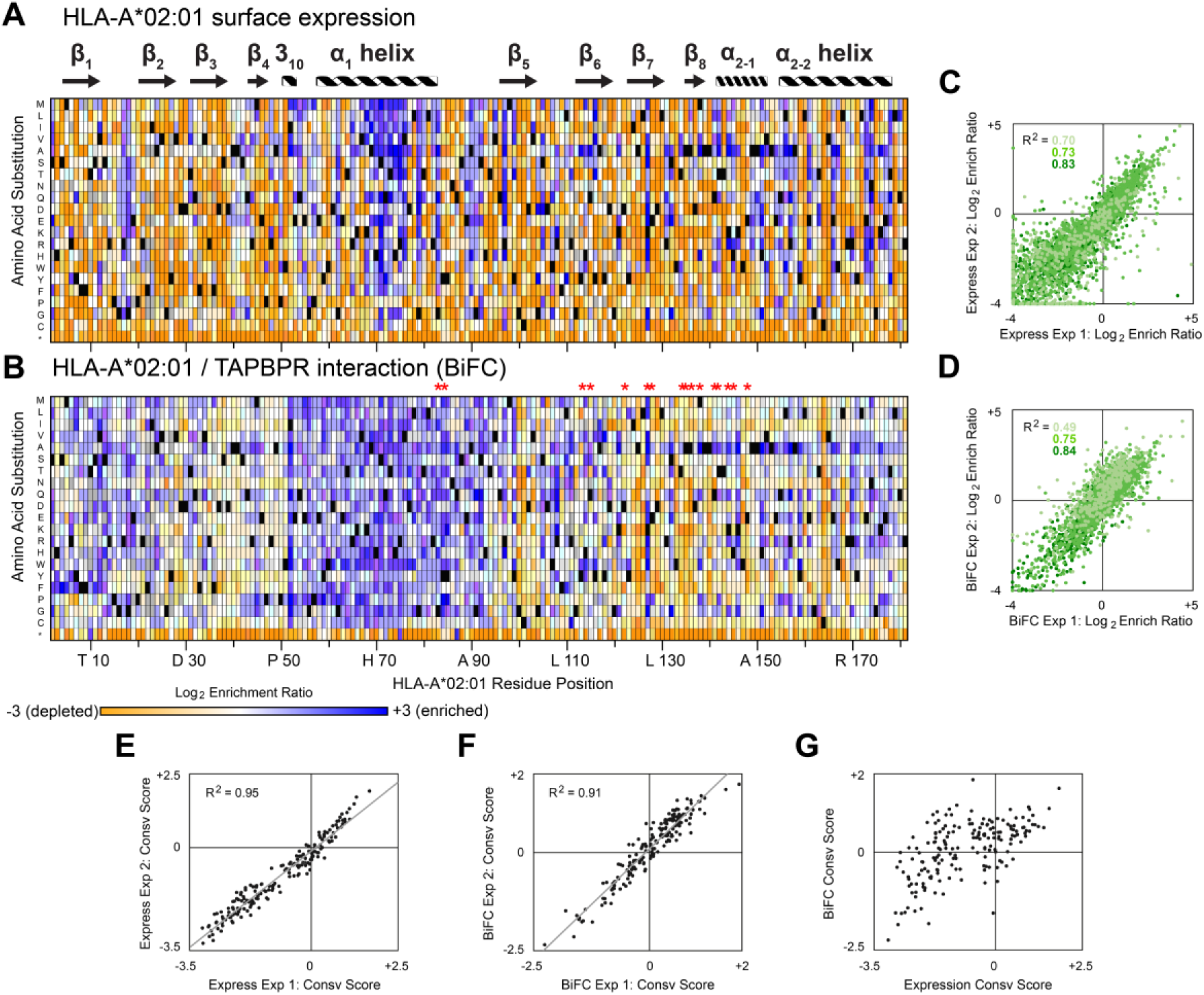
Mutational landscapes of HLA-A*02:01 reveal relaxed sequence constraints for TAPBPR binding. **(A)** A site-saturation mutagenesis library of HLA-A*02:01 was selected by FACS for high surface expression. Residue position is on the horizontal axis, and amino acid substitutions are on the vertical axis (*, stop codon). Log_2_ enrichment ratios are plotted from ≤ −3 (i.e. depleted/deleterious, orange) to ≥ +3 (enriched, dark blue). Mutations missing in the naive library (frequency ≤ 2×10^−6^) are grey, wild-type amino acids are black. **(B)** The VN-fused HLA-A*02:01 library was selected by FACS for high BiFC in cells expressing VC-fused TAPBPR-TM. Colored as in panel B. Red asterisks (*) denote HLA-A*02:01 residues in direct contact with TAPBPR based on the homologous H2-D^d^/TAPBPR X-ray structure (PDB ID 5WER). **(C and D)** Correlation plots showing the agreement of log_2_ enrichment ratios for all mutations between two independent selection experiments for (C) HLA-A*02:01 surface expression or (D) HLA-A2/TAPBPR-TM BiFC. Mutations are binned from low (2×10^−6^ to 2×10^−5^; pale green) to medium (2×10^−5^ to 2×10^−4^; medium green) to high (≥ 2×10^−4^; dark green) frequency in the naive library. **(E and F)** Agreement between residue conservation scores (calculated from the mean of the log_2_ enrichment ratios for all non-stop mutations at a given position) from independent replicate selections for (E) HLA-A2 surface expression or (F) HLA-A*02:01/TAPBPR-TM BiFC. Conserved residues have negative scores. **(G)** Comparison of residue conservation scores between selections of the HLA-A*02:01 library for surface expression versus BiFC with TAPBPR-TM, showing that the HLA-A*02:01 sequence is more tolerant of mutations for TAPBPR interactions.

**Fig. S5.**
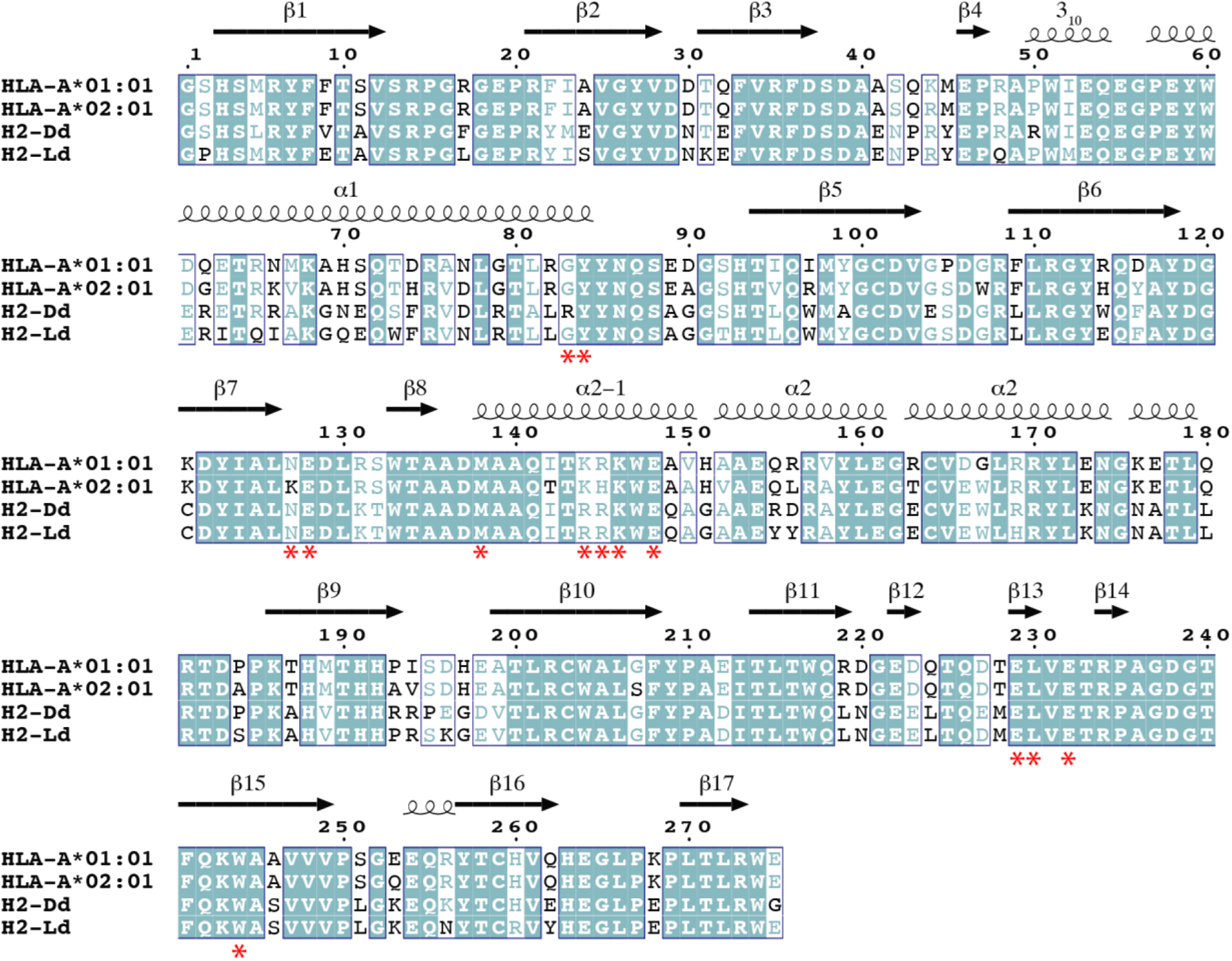
Protein sequence alignment of MHC-I molecules characterized by NMR in this study. Sequence alignment of the luminal domains of H2-D^d^ (UniProtKB/Swiss-Prot: P01900), H2-L^d^ (UniProtKB/Swiss-Prot: P01897), HLA-A*01:01 (UniProtKB/Swiss-Prot: P30443), and HLA-A*02:01 (UniProtKB/Swiss-Prot: P01892) performed using ClustalOmega (*87*) and processed with ESPript (*88*). Secondary structure of the heavy chain from H2-D^d^ (PDB ID 3ECB) is provided for reference. Residues in blue boxes are conserved. The red asterisk (*) denotes heavy chain residues that are in direct contact with TAPBPR based on the structure of H2-D^d^ and TAPBPR (PDB 5WER) as calculated with the software Protein Interfaces, Surfaces and Assemblies (PISA) (*89*).

**Fig. S6.**
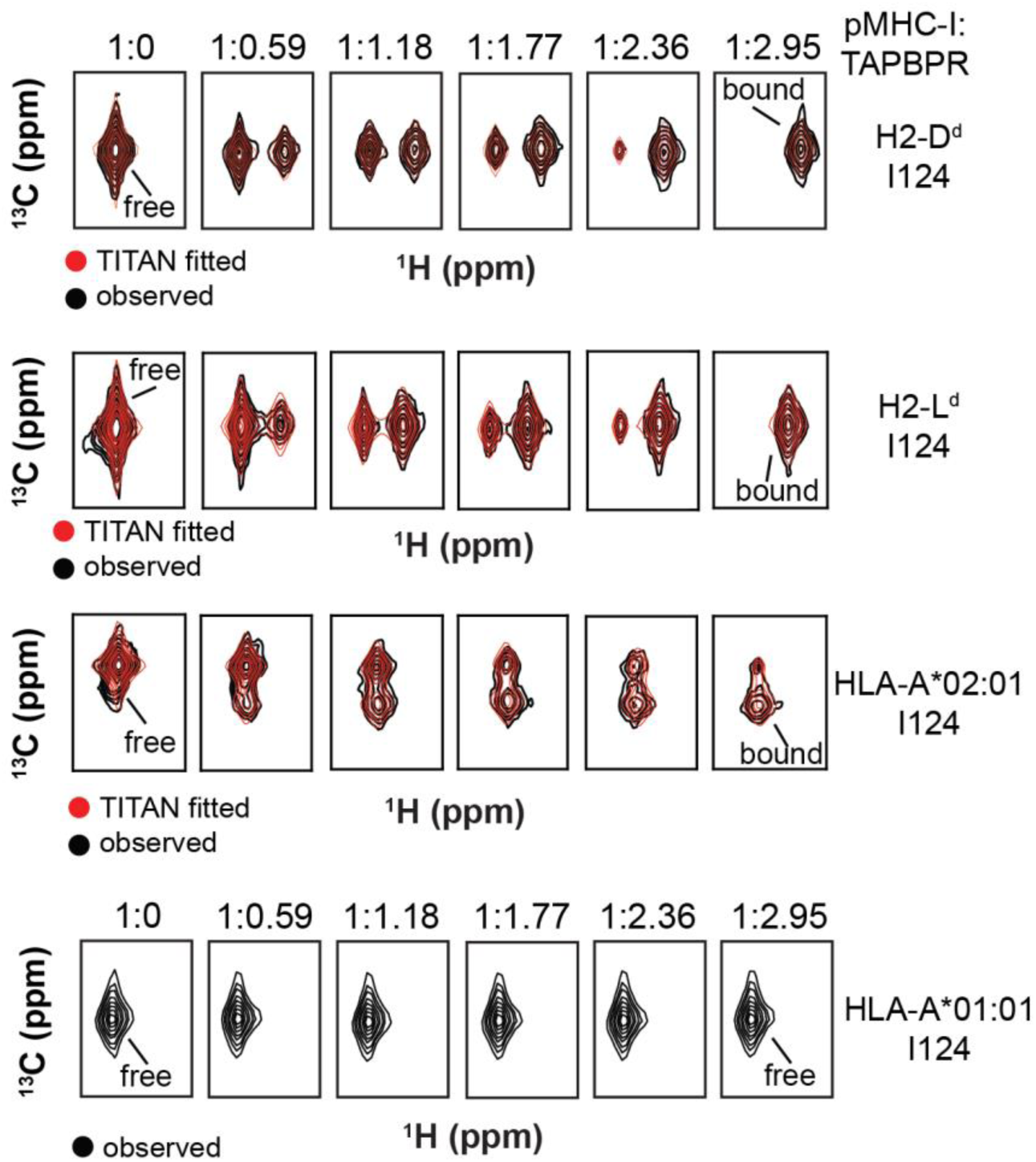
Representative titrations of pMHC-I with TAPBPR. NMR line shape analysis, performed in TITAN (*84*), of the ^13^C_δ1_ methyl resonance corresponding to residue I124 for H2-D^d^, H2-L^d^, HLA-A*02:01 and HLA-A*01:01. The experimental NMR line shapes are colored black, and the TITAN fits are colored red. Snapshots from the 2D ^1^H-^13^C SOFAST HMQC experiment for the NMR peak of I124 δ1 methyl are shown throughout the titration at different molar ratios of the pMHC-I to TAPBPR (further details in the Materials & Methods section). The initial concentration of the pMHC-I was 105 μM (1:0 titration point) and experiments were recorded at 25°C at 800 MHz. The NMR peaks of the free (pMHC-I) and bound (pMHC-I/TAPBPR) states are noted. HLA-A*01:01 did not exhibit TAPBPR binding and thus was not fitted.

**Fig. S7.**
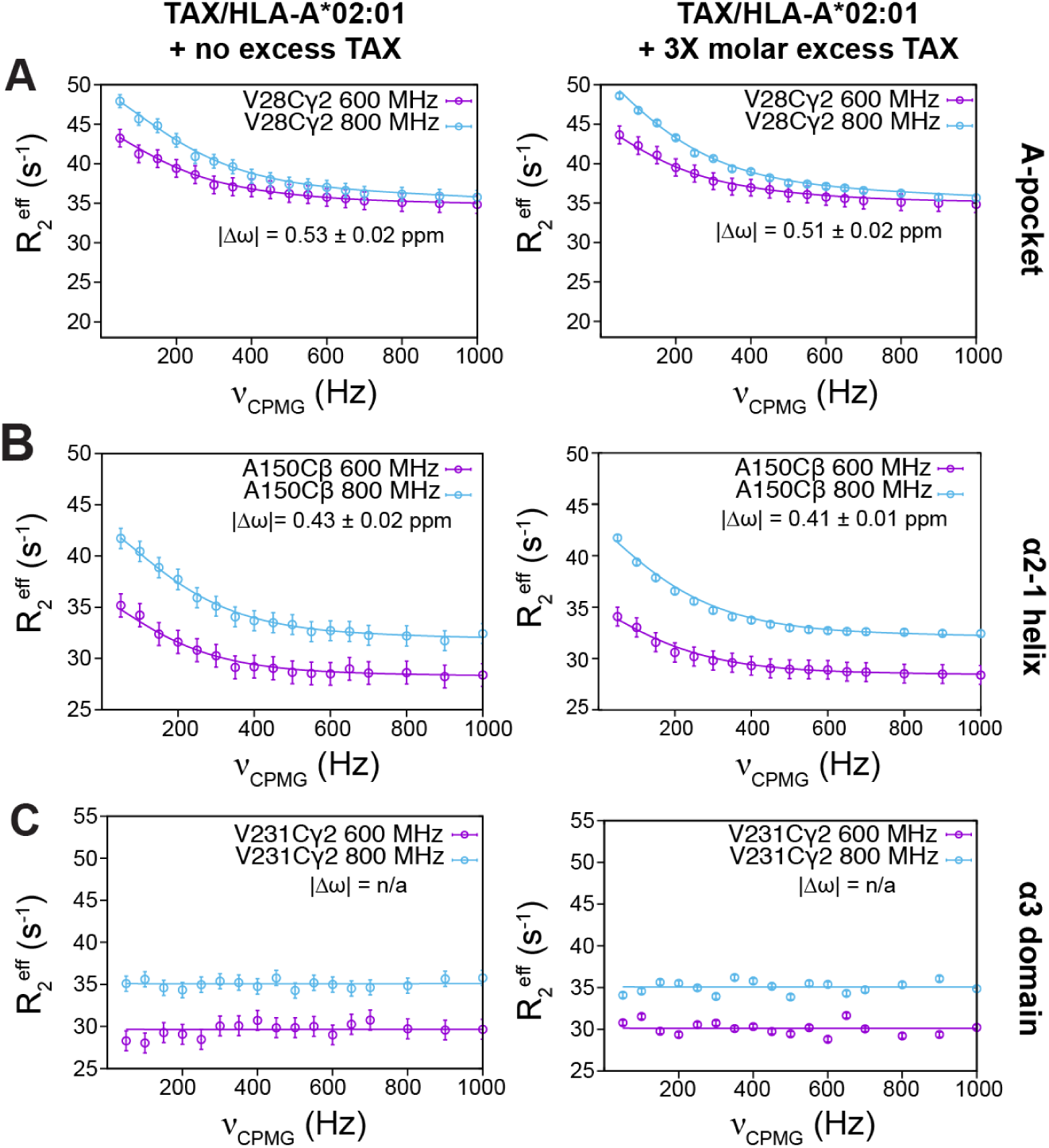
Comparison of ^13^C-SQ CPMG relaxation dispersion profiles recorded for free TAX/HLA-A*02:01/hβ2m at 1:1 and 3:1 peptide:MHC-I ratios. Profiles of ^13^C-SQ CPMG relaxation dispersion experiments (*53*) recorded at 25°C (800 MHz – blue; 600 MHz – purple) on AILV methyl labeled (at the heavy chain) pMHC-I. Methyl groups of each heavy chain in specified regions, including the **(A)** A-pocket, **(B)** α_2-1_ helix, and **(C)** α_3_ domain, exhibit allele-specific conformational exchange. Flat profiles indicate no observable μs-ms dynamics, compared to profiles exhibiting “curve” behavior, which are undergoing significant conformational exchange. The effective transverse relaxation rate (R_2_^eff^, s^−1^, y-axis) is shown as a function of the CPMG pulse frequency (ν_CPMG_, Hz, x-axis). For each pMHC-I molecule the global fit of the relaxation dispersion curves performed in CATIA (http://www.biochem.ucl.ac.uk/hansen/catia/) are shown. Upper and lower error bars of R_2_^eff^ were determined based on the spectral noise. The fitted |Δω| values are noted in each panel. CATIA fitted parameters are *k_ex_* = 1102 ± 16 s^−1^ and *p_B_* = 5.21 ± 0.02% (no excess peptide) and *k_ex_* = 1126 ± 18 s^−1^ and *p_B_* = 5.56 ± 0.02% (three-fold molar excess TAX peptide).

**Fig. S8.**
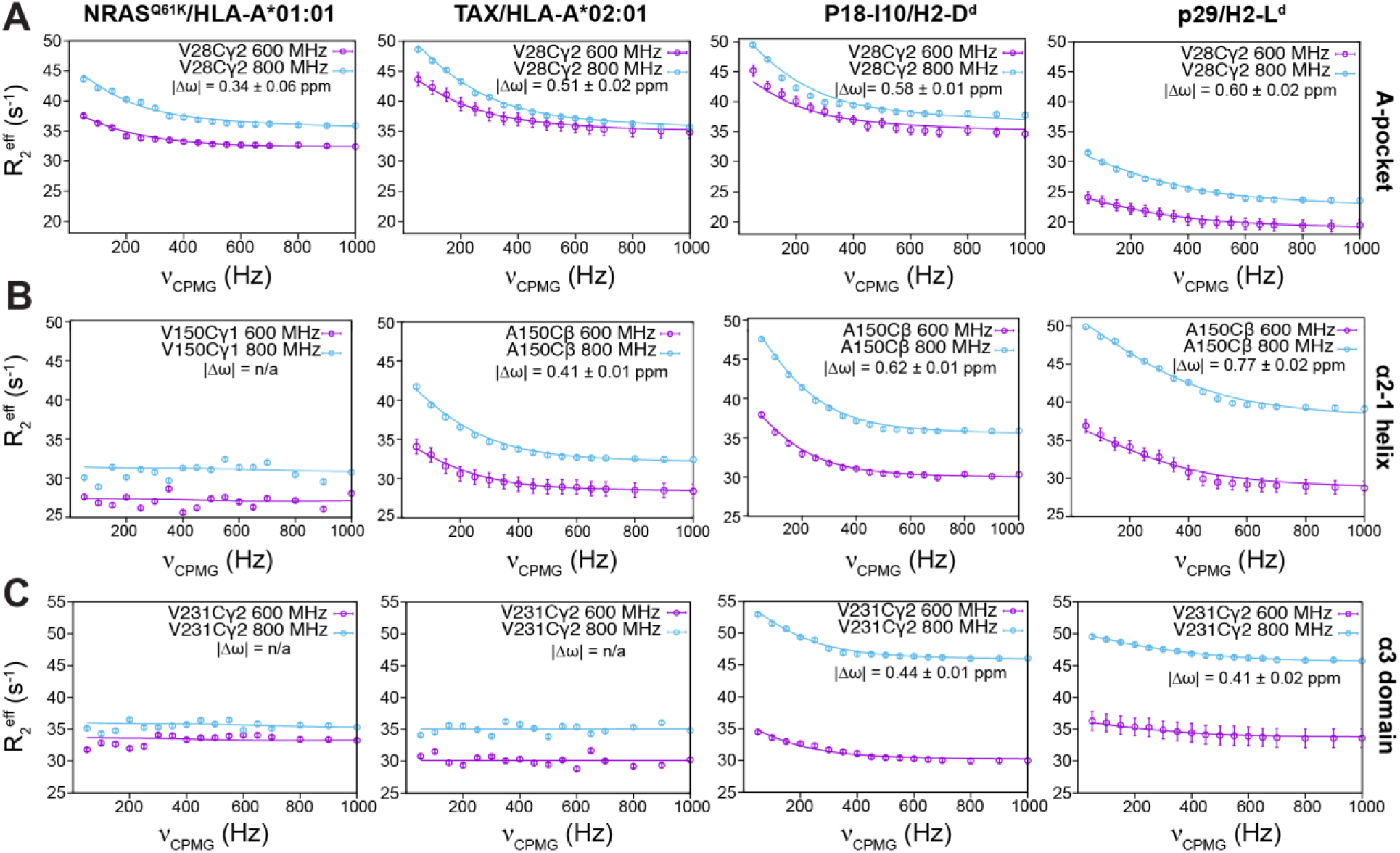
Representative ^13^C-SQ CPMG relaxation dispersion curves highlighting site-specific conformational exchange in unchaperoned pMHC-I molecules. Profiles of ^13^C-SQ CPMG relaxation dispersion experiments (*53*) recorded at 25°C (800 MHz – blue; 600 MHz – purple) on AILV methyl labeled (at the heavy chain) pMHC-I. Methyl groups of each heavy chain in specified regions, including the **(A)** A-pocket, **(B)** α_2-1_ helix, and **(C)** α_3_ domain, exhibit allele-specific conformational exchange. Flat profiles indicate no observable μs-ms dynamics, compared to profiles exhibiting “curve” behavior, which are undergoing significant conformational exchange. The effective transverse relaxation rate (R_2_^eff^, s^−1^, y-axis) is shown as a function of the CPMG pulse frequency (ν_CPMG_, Hz, x-axis). For each pMHC-I molecule the global fit of the relaxation dispersion curves performed in CATIA (http://www.biochem.ucl.ac.uk/hansen/catia/) are shown. Upper and lower error bars of R_2_^eff^ were determined based on the spectral noise. The fitted |Δω| values are noted in each panel and can also be found in Table S2.

**Fig. S9.**
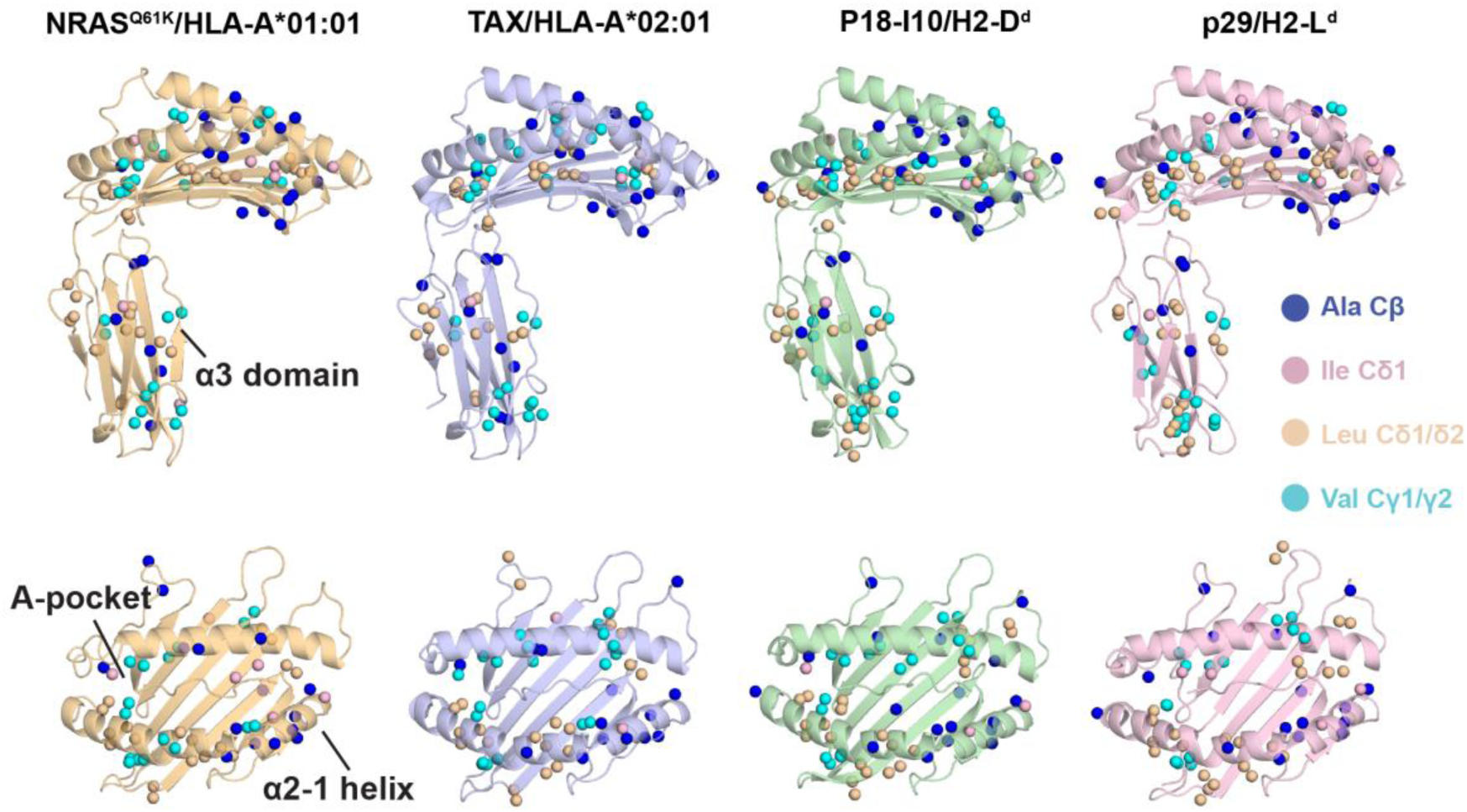
Distribution of assigned AILV methyl probes on pMHC-I structures. Views of the pMHC-I molecules used in this study are shown from the side (upper images) and top of the groove (lower images). Regions of interest (A-pocket, α_2-1_ helix and α_3_ domain) are noted. Methyl groups with assigned Ala C_β_, Ile C_δ1_, Leu C_δ1_/C_δ2_, and Val C_γ1_/C_γ2_ methyl resonances are shown as spheres with colors denoted in the legend on the right. The PDB IDs of the 4 alleles used are: HLA-A*01:01 (PDB ID 6MPP, light orange), HLA-A*02:01 (PDB ID 1DUZ, light blue), H2-D^d^ (PDB ID 3ECB, light green) and H2-L^d^ (PDB ID 1LD9, light pink).

**Fig. S10:**
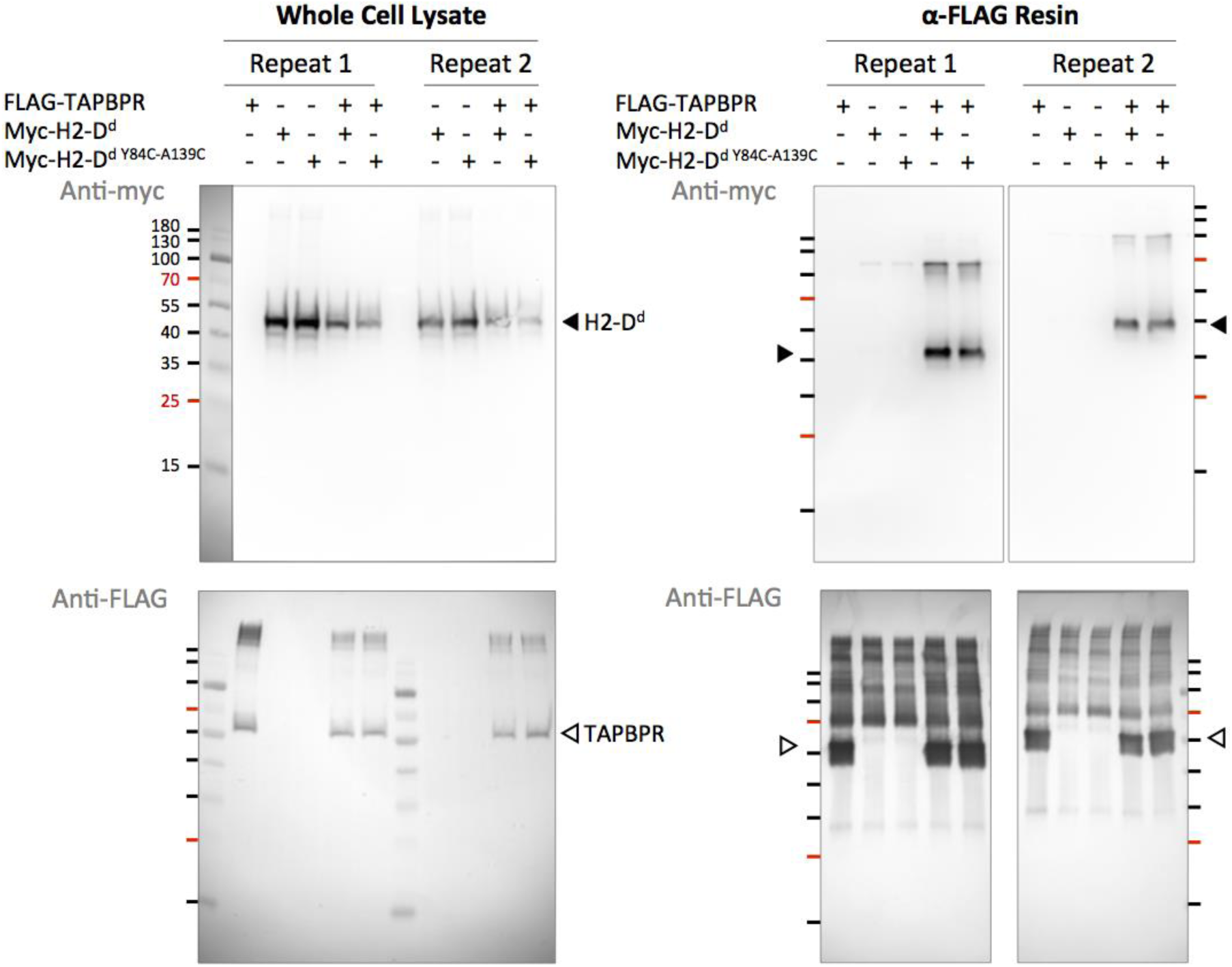
Introduction of cysteines at positions 84 and 139 of H2-D^d^ does not prevent association with TAPBPR in cells. Human Expi293F cells were transfected with FLAG-tagged TAPBPR and/or myc-tagged H2-D^d^. Total expression in whole cell lysate is shown by anti-myc (upper) and anti-FLAG (lower) immunoblots at left. TAPBPR complexes were immunoprecipitated with anti-FLAG resin, and after consideration of slight differences in total expression, H2-D^d^ and H2-D^d Y84C-A139C^ coprecipitate with TAPBPR at similar levels based on immunoblots at right. Sizes of molecular weight markers are indicated in kDa. Filled arrowheads indicate H2-D^d^, open arrowheads indicate TAPBPR.

## Supporting Tables

**Table S1.** Thermal stability of different pMHC-I molecules does not correlate with affinity for TAPBPR in vitro. Melting temperatures (T_m_, °C) obtained from differential scanning fluorimetry experiments (*45*) are reported for each pMHC-I. Standard errors obtained from triplicate measurements are shown (n = 3). T_m_ values were fit using a Boltzmann sigmoidal function in GraphPad Prism v7.

| pMHC-I | T<sub>m</sub> (°C) |
| --- | --- |
| P18-I10/H2-D<sup>d</sup>/hβ2m | 55.0 ± 0.1 |
| P18-I10/H2-D<sup>d</sup> Y84C-A139C/hβ2m | 53.0 ± 0.3 |
| p29/H2-L<sup>d</sup>/hβ2m | 56.3 ± 0.2 |
| TAX/HLA-A*02:01/hβ2m | 63.5 ± 0.1 |
| NRAS<sup>Q61K</sup>/HLA-A*01:01/hβ2m | 51.4 ± 0.1 |

**Table S2.**
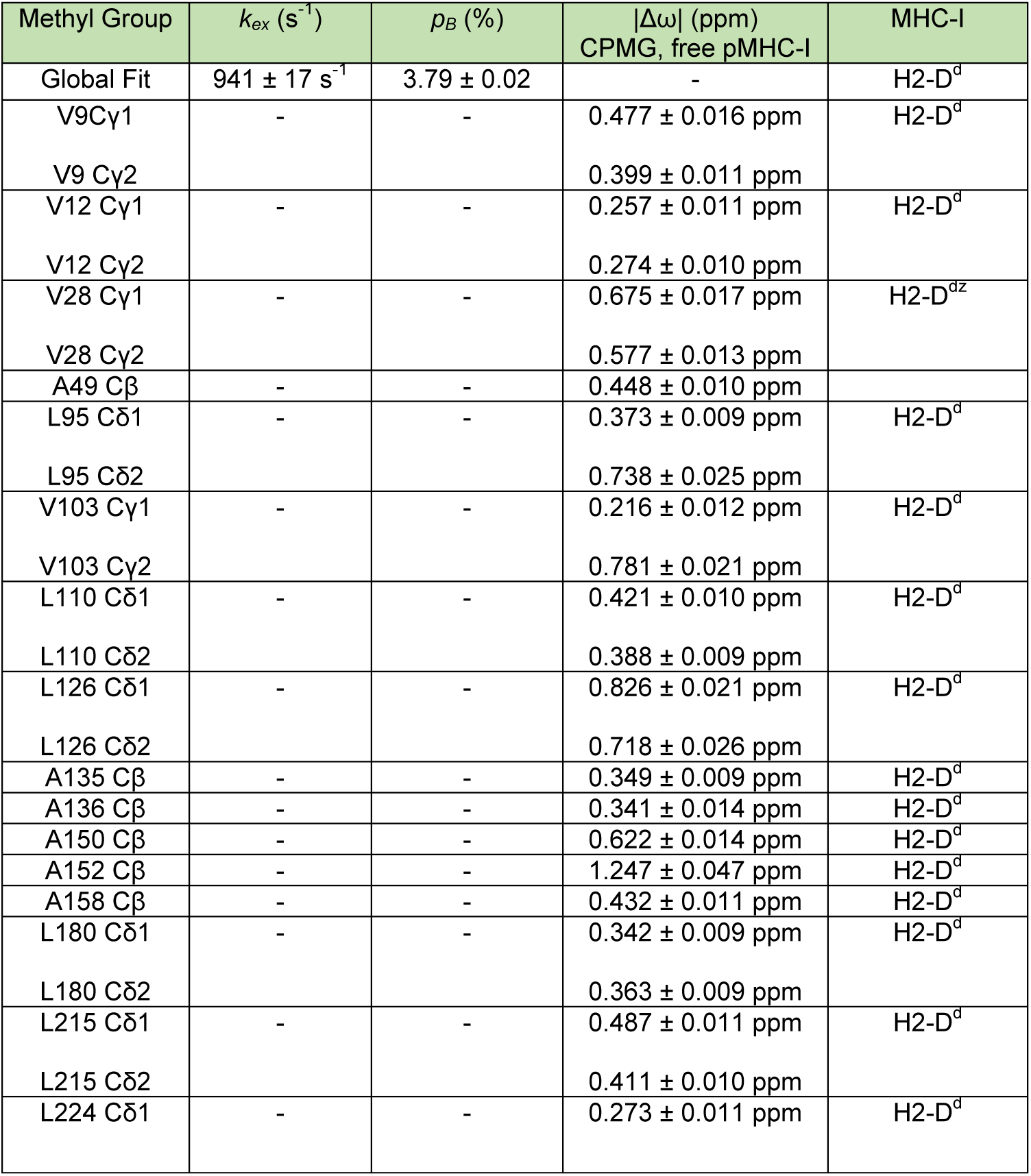
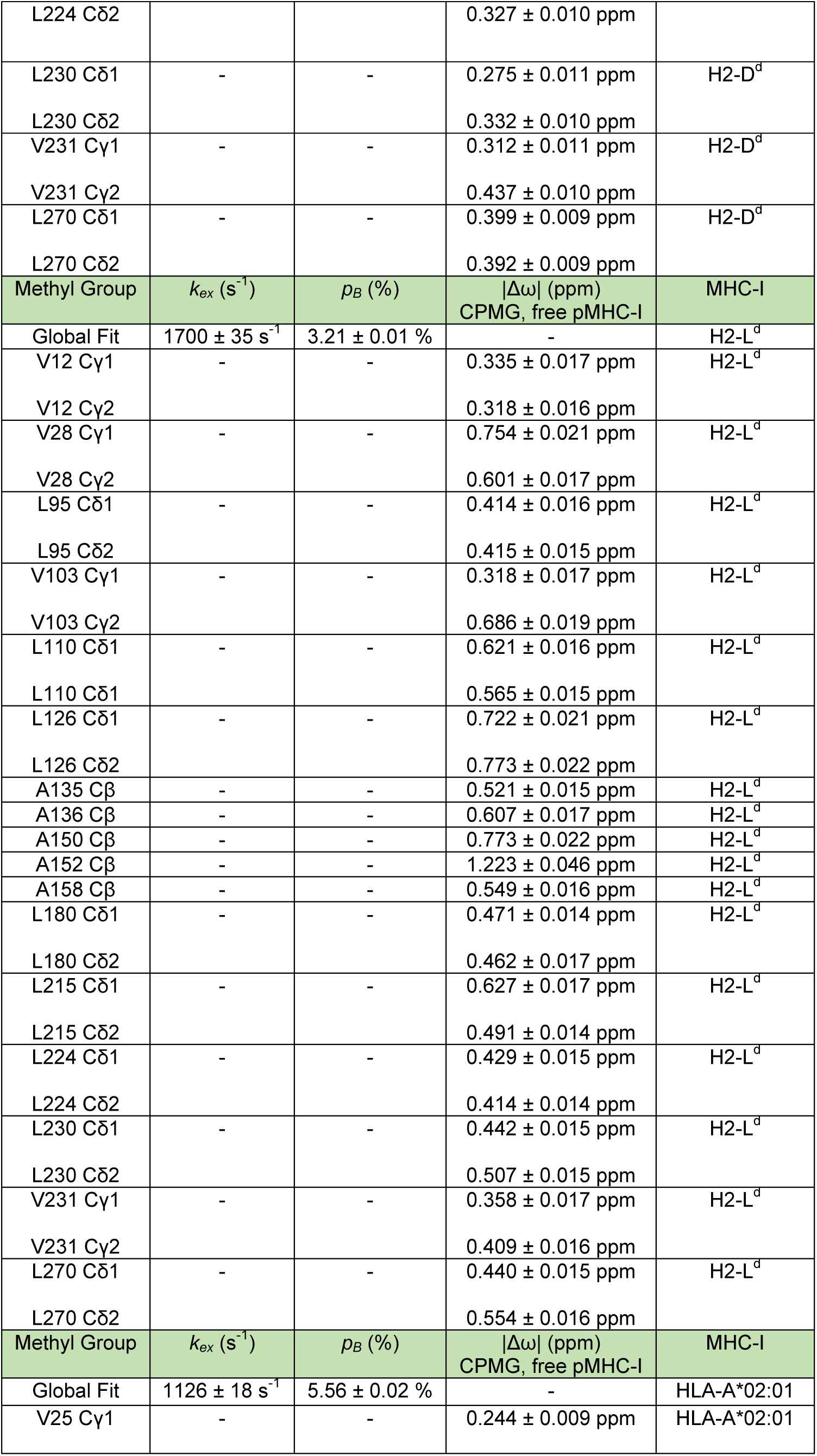
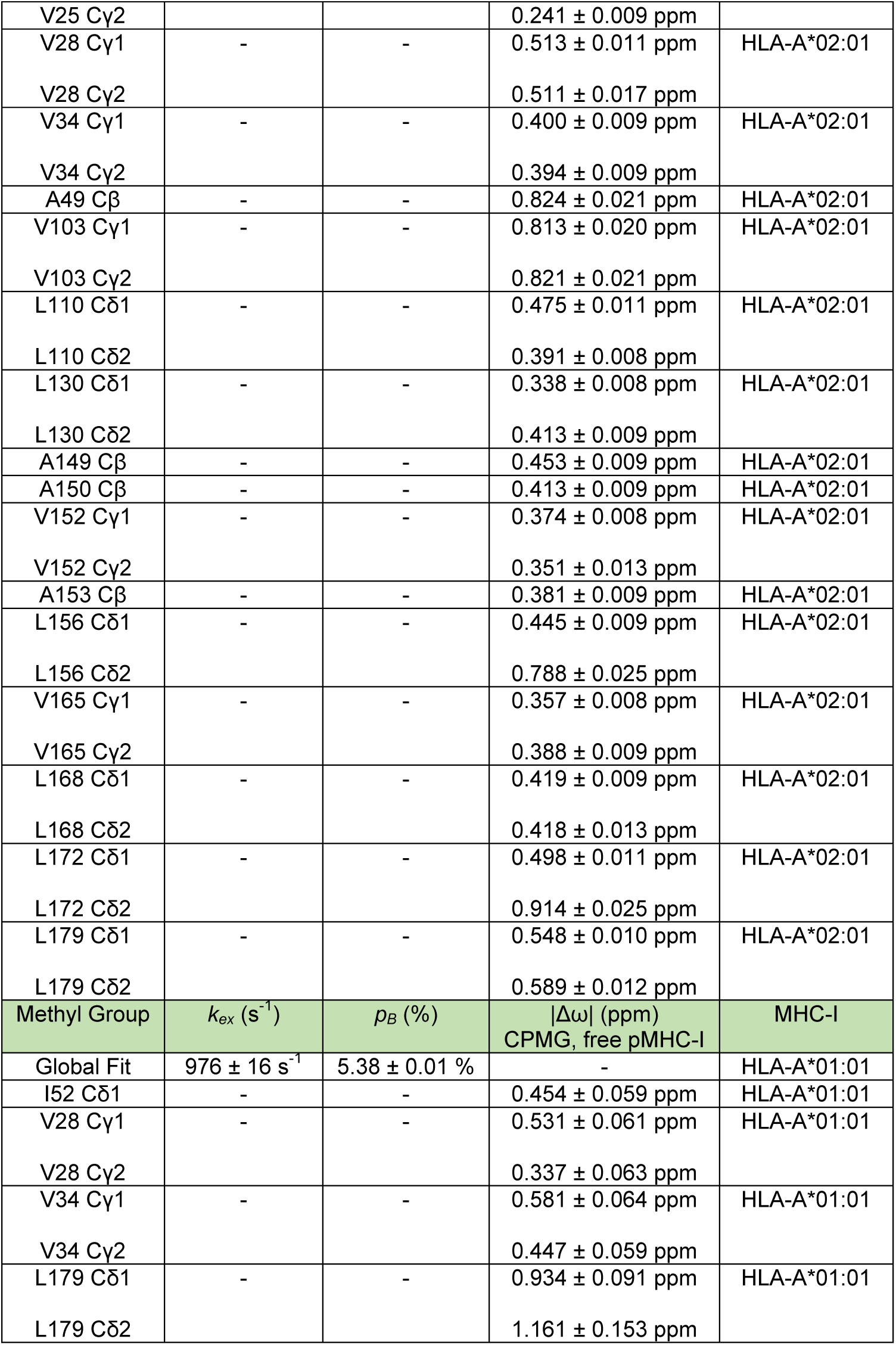
Summary of parameters obtained from a global fit of ^13^C-SQ CPMG relaxation dispersion curves, performed in CATIA of different pMHC-I molecules at ^1^H NMR fields of 600 and 800 MHz at 25°C.

**Table S3:**
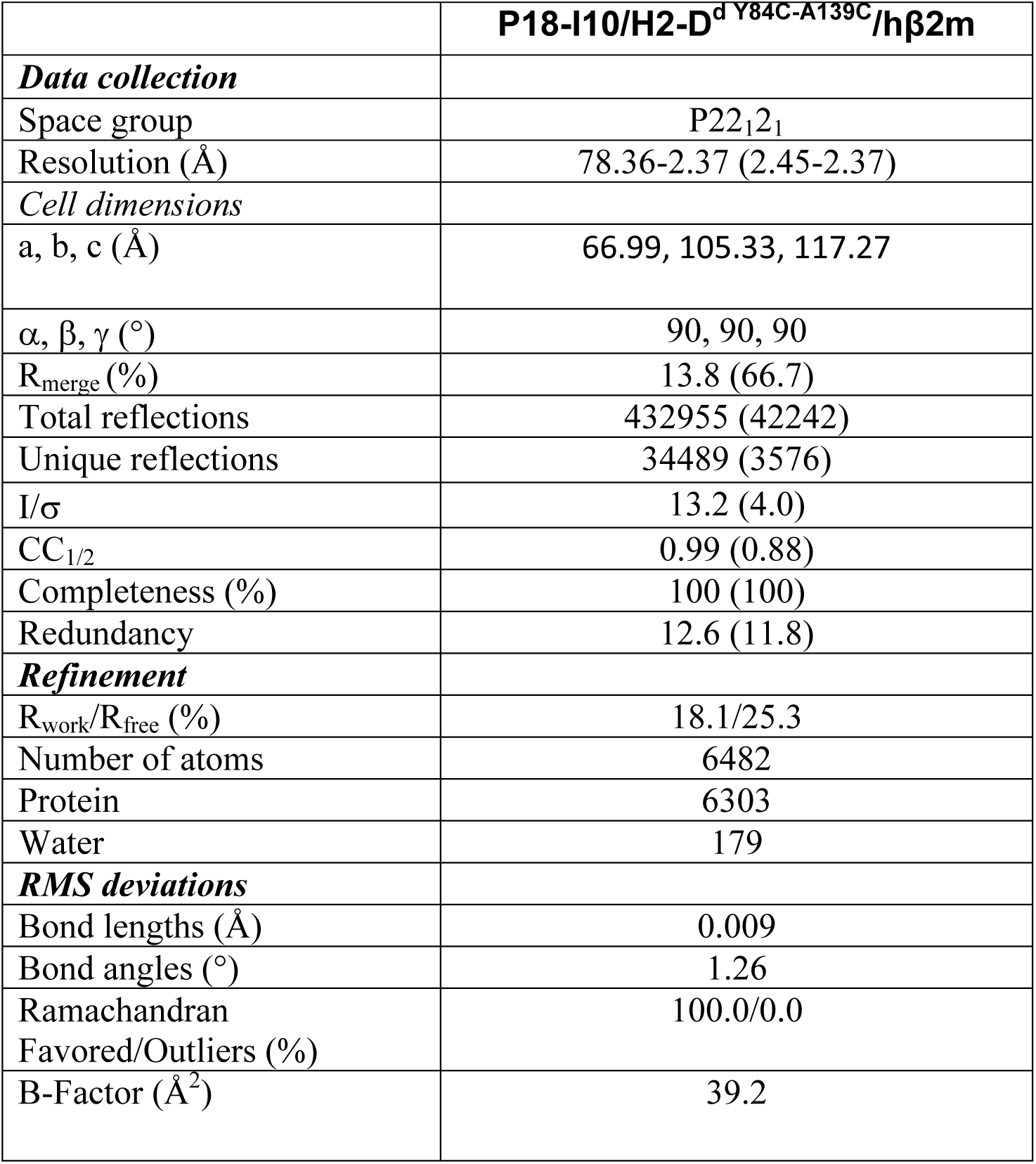
X-ray crystallography data collection and refinement statistics for the P18-I10/H2-D^d Y84C-A139C^/hβ2m complex. Values in parentheses in the right column correspond to the highest resolution shell. PDB ID 6NPR.

